# Proteostasis perturbation destabilizes respiratory complex assembly-intermediates via aggregation of subunits

**DOI:** 10.1101/312793

**Authors:** Shivali Rawat, Valpadashi Anusha, Manoranjan Jha, K Sreedurgalakshmi, Anamika Ghosh, Debabani Ganguly, Swasti Raychaudhuri

## Abstract

Proteostasis is maintained by optimum expression, folding, transport, and clearance of proteins. Deregulation of any of these processes triggers widespread protein aggregation and loss of function. Here, we perturbed proteostasis by blocking proteasome-mediated protein degradation and investigated proteome partitioning from soluble to insoluble fraction. Aggregation of Respiratory Chain Complex (RCC) subunits highlights the early destabilization event as revealed by proteome redistribution. Sequence analyses followed by microscopy suggest that low complexity regions at the N-terminus are capable to facilitate aggregation of RCC subunits. As a result, respiratory complex assembly process is impaired due to destabilization of sub-complexes marking the onset of mitochondrial dysfunction and ROS accumulation. Redistribution of Histone proteins and their modifications indicated reprogramming of transcription as adaptive response. Together, we demonstrate susceptibility of RCC subunits to aggregation under multiple proteotoxic stresses providing an explanation for the simultaneous deregulation of proteostasis and bioenergetics in age-related degenerative conditions.

## INTRODUCTION

Any physicochemical stress that may perturb protein-conformation is capable to trigger misfolding and insolubility. Heat stress unfolds the secondary structures, exposes the hydrophobic stretches of proteins and results in the formation of either reversible or amyloidogenic aggregates (Audas et al., 2016; Wallace et al., 2015). Similarly, imbalance of protein homeostasis (proteostasis) network can also promote insolubility as the protein-stabilizing factors are gradually sequestered by the increasing load of misfolded proteins (Vabulas et al., 2010). However, the identity of the destabilized proteins and the magnitude of instability may differ depending on the nature of proteostasis stresses and cellular growth conditions. Moreover, perturbing the same node of proteostasis network may trigger conformational collapse and insolubility of different groups of proteins in condition-specific manner. For example, members of Hsp70 chaperone family normally protect the hydrophobic stretches of the newly synthesized proteins from unproductive folding and aggregation. However, limited availability of the same chaperones results in aggregation of mature but unfolded proteins by decelerating refolding in presence of proteotoxic stresses (Vabulas et al., 2010). Predominantly, the misfolded proteins are unable to perform their normal functions and partition into the insoluble fraction. However, proteome-destabilization events may not be always deleterious. Multiple studies using cell culture and aging models have suggested that transient phase-separation of proteins is a part of cellular defence in proteotoxic conditions (Audas et al., 2016; Wallace et al., 2015; Walther et al., 2015).

In this study, we induced proteome instability in mammalian cells by perturbing the turnover of proteins. In mitotic cells, majority of the cellular proteins are stable enough to be transferred to the daughter cells after division. However, several signaling proteins like cell cycle enzymes, transcription factors etc., which constitute about 5% of the cellular proteome, have significantly lesser half-life and are rapidly degraded after transient expression and function. Thus, turnover profile of certain proteins is fairly dynamic depending on the cell cycle stage, sub-cellular localization or environmental fluctuations (Goldberg, 2003; Muratani and Tansey, 2003). A block in the timely clearance of these proteins by the ubiquitin-proteasome system (UPS), autophagy or by organellar proteases not only triggers wrong signalling events but also surges a misfolded protein load that overwhelms the overall proteostasis-capacity and creates a competition among the chaperone-dependent proteins for productive folding. The destabilized proteome precipitates over time, results in accumulative proteotoxicity and cell death (Tanaka and Matsuda, 2014; Vilchez et al., 2014).

Large-scale proteomic studies investigating the impact of proteasome-inhibition on the cellular proteome have already identified several short-lived proteasome substrates and delineated compartment-specific protein degradation mechanisms (Bieler et al., 2012; Bieler et al., 2009; Larance et al., 2013; Wilde et al., 2011). Along with the polyubiquitinated proteins, multiple chaperones and protein-degradation machinery components have been identified in the insoluble fraction of these cells suggesting a cellular attempt to clear the damaged proteins. Despite these highthroughput attempts, it remained to be investigated whether widespread insolubility of proteins due to proteasome-inhibition is stochastic in nature or organized and, how do these aggregation events contribute to cellular protection or toxicity. To address, we captured a snapshot of proteome-reorganization events in Neuro2a cells after short-term proteasome inhibition. Partitioning of mitochondrial proteins, especially the respiratory chain complex (RCC) subunits from soluble to insoluble fraction was reflected as a major perturbation in the proteome. Many of these nuclear-encoded RCC subunits were found to contain N-terminal low complexity regions (LCRs) that contributed to the aggregation events. Complexome profiling experiments revealed that destabilization of small assembly intermediates composed of these aggregation-prone RCC subunits earmarks the onset of long-term respiration stress. Redistribution of multiple histone-proteins across the proteome fractions, increased load of H3K4 trimethylation and induction of hsp-transcription were identified as early-adaptive responses in the pursuit of re-establishing the proteome-equilibrium.

## RESULTS

### Destabilization of chaperone-dependent proteins in Neuro2a cells by short-term proteasome-inhibition

To study proteome partitioning from soluble to insoluble fraction, we lysed Neuro2a cells using a mild lysis buffer containing 1% NP-40 and 0.25% deoxycholate (Gupta et al., 2011; Walther et al., 2015). Although most of the cellular proteins including components of various organelles were soluble in this buffer **(Figure S1A)**, we always obtained a pellet after centrifuging the lysate. We dissolved the pellet in 2% SDS containing buffer and found cytoskeletal protein β-actin and chromatin associated protein Histone H3 as significant constituents of this insoluble fraction. Endoplasmic reticulum chaperone Grp78 was found to be associated with the insoluble proteins although mitochondrial Hsp60 and cytosolic Hsp70 were not abundant in this fraction under normal conditions **(Figure S1A)**. We also dissolved the cells directly in SDS to prepare the total fraction that contain both NP-40 soluble and insoluble proteins.

Next, we treated mouse neuroblastoma cell-line Neuro2a with two different concentrations (2.5 and 5 μM) of a reversible cell-permeable proteasome inhibitor MG132 (Kisselev and Goldberg, 2001) to select a condition that destabilizes the proteome but do not trigger cell death. Treatment for 8 hr was sufficient to block proteasome activity almost completely with increased load of ubiquitinated proteins but no loss of cell viability (**Figure 1A and S1B-D**). Then, we transfected cells with a reporter-construct expressing FlucDM-EGFP; Firefly luciferase with double mutations (FlucDM), a conformationally unstable, chaperone dependent protein that acts as proteostasis-stress sensor (Gupta et al., 2011). At optimum growth conditions, FlucDM-EGFP was found to be diffusely distributed throughout with small dispersed IBs in few cells. Treatment with MG132 (2.5 and 5 μM) resulted in larger IBs by 8 hr (**Figure 1B**). Western blot of the cell-fractions confirmed increased load of FlucDM-EGFP in total fraction due to reduced degradation in presence of MG132 and subsequent precipitation to insoluble fraction. However, no change in the protein level was observed in soluble fraction (**Figure S1E**). Simultaneous drop in luminescence-activity suggested a dose-dependent loss-of-activity of FlucDM-EGFP due to increased instability (**Figure 1C**). Hsp70 protein-load was also increased in the total and insoluble fraction of these cells, but not in soluble fraction, suggesting its association with destabilized FlucDM-EGFP (**Figure S1F**).

**Figure 1:**
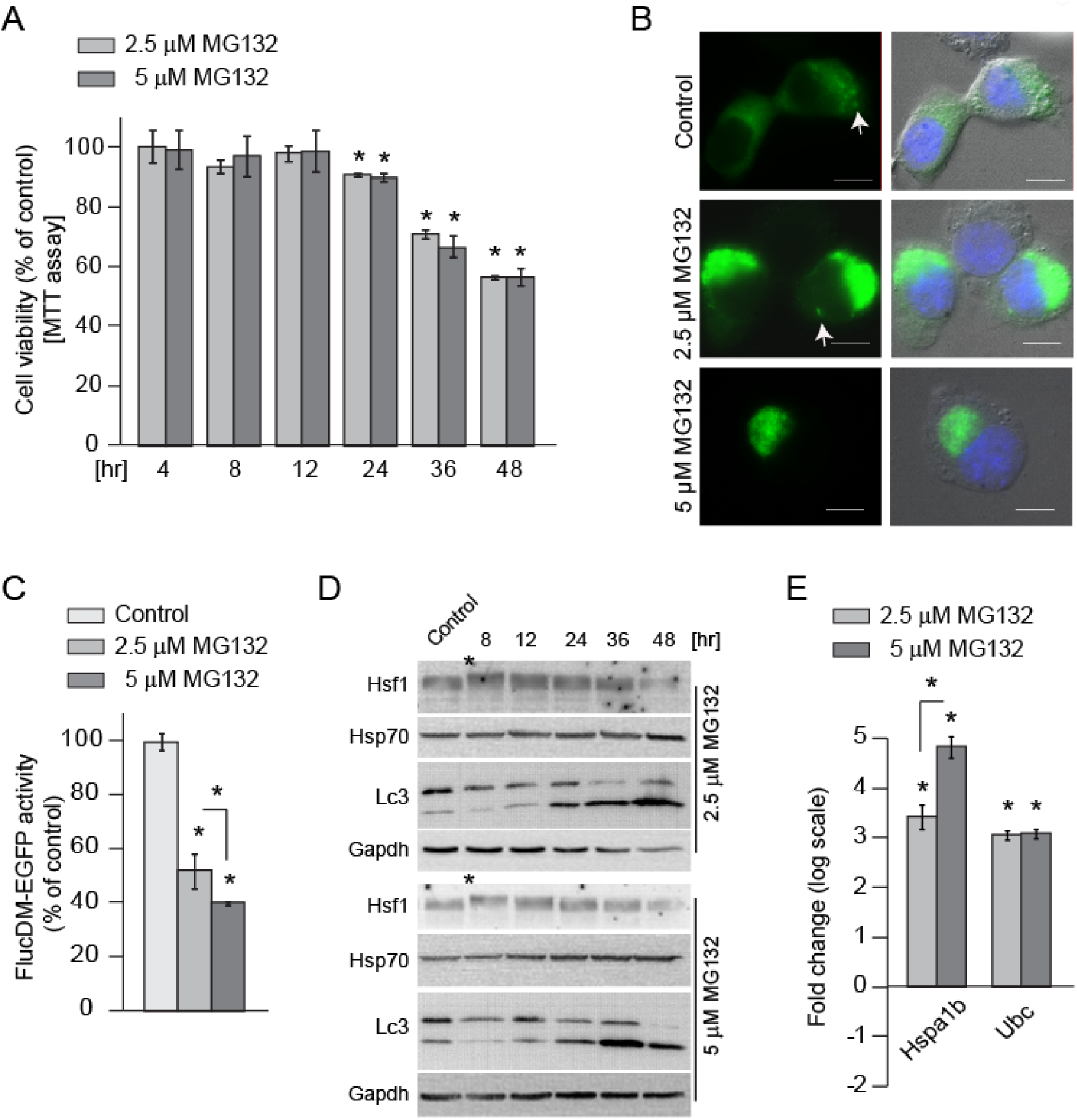
Destabilization of chaperone-dependent proteins in Neuro2a cells by short-term proteasome-inhibition. (A) Cell viability. Neuro2a cells were incubated with MG132 and cell viability was estimated by MTT assay. Control: DMSO treated cells. Error bars indicate SDs from at least three independent experiments. ^*^ indicates p<0.05 by Student’s t-test. (B) Fluorescence micrographs showing IBs formed by over-expressed FlucDM-EGFP in MG132-treated Neuro2a cells for 8 hr. Nucleus is stained by DAPI. Arrows indicate smaller IBs. Control: DMSO treated cells. Scale-bar - 10 μm. (C) FlucDM activity assay. Luciferase activity was measured in FlucDM-EGFP transfected cells as described in Methods. Control: DMSO treated cells. Error bars indicate SDs from at least three independent experiments. ^*^ indicates p<0.05 by Student’s t-test. (D) Hsp70, Hsf1 and Lc3 protein-levels in MG132-treated cells. Neuro2a cells were treated with MG132 for different time lengths, soluble extracts were prepared and immunoblot was performed with anti-Hsp70, anti-Hsf1, anti-Lc3 and anti-GAPDH. ^*^ indicates upshifted band of Hsf1. Extract of DMSO treated cells was used as control. Gapdh served as loading control. (E) Hsp70 and ubiquitin mRNA-levels in MG132-treated cells for 8 hr as determined by real-time PCR. Fold change was normalized against mRNA levels in DMSO treated cells. Error bars indicate SDs from at least three independent experiments. ^*^ indicates p<0.05 by Student’s t-test. See also **Figure S1**.

MG132 treatment is known to trigger defence mechanisms like Heat shock factor 1 (Hsf1) mediated cytosolic stress response (Heat shock response; HSR) (Westerheide and Morimoto, 2005). Hsf1normally remain sequestered in the cytoplasm as a complex with chaperones like Hsp90 and Hsp70. In presence of protein-destabilization events, these chaperones dissociate from this complex to protect the misfolded cytosolic proteins resulting in oligomerization and post-translational modifications of free monomeric Hsf1 towards trans-activation (Raychaudhuri et al., 2014). We found trans-activated Hsf1 by 8 hr treatment of MG132 as observed by the upshifted phosphorylated band in the western blot suggesting presence of misfolded proteins in the cytosol that are capable to dissociate the Hsf1-chaperone complex (**Figure 1D)**. Simultaneous increase in mRNA-levels of hsp70 (hspa1b) and Ubiquitin (ubc) genes supported upregulation of Hsf1-mediated transcription (**Figure 1E**). The upshifted Hsf1 band started migrating to the original position by 24 hr illustrating a reversion of the trans-activation mechanism (**Figure 1D**).

Often, autophagy has been reported to be induced in proteasome-inhibited cells as an adaptive mechanism for misfolded protein clearance (Bao et al., 2016; Barth et al., 2010). In our experiments, an increased cleavage of LC3 was observed indicating activation of autophagy only by 24 hr; however, uncleaved version was prominent at 8 hr **(Figure 1D).** To note, a gradual reduction in cell viability was also observed by 24 hr implying that the adaptive rescue mechanisms were not sufficient to protect the cells during long-term proteasome inhibition (**Figure 1A and S1B**). Thus, the 8 hr of MG132 treatment represented a timepoint when the partitioning of the conformationally unstable proteins to the non-functional insoluble fraction just started contributing to proteotoxicity; however, cellular functions are still unaffected and adaptive mechanisms are waking up sensing the minute equilibrium shift in the proteome.

### Proteome reorganization in response to proteasome inhibition

In order to probe the proteome partitioning events in MG132 treated cells by 8 hr, we quantified 1986 proteins in total fraction, 1329 proteins in soluble fraction and 838 proteins in insoluble fraction by SILAC-based quantitative mass spectrometry (**Figure 2A and S2A; Table S1**). The proteins in total and soluble fraction were components of multiple cell-compartments and participating in various cellular activities. The 838 proteins identified in insoluble fraction were physicochemically distinct with basic isoelectric point (pI), more low complexity regions (LCRs) and +vely charged residues and highly abundant as per iBAQ analysis **(Figure S2B)**. Contrary to the general perception that membrane association causes proteins to precipitate (Monsellier et al., 2008), relatively small number of transmembrane helix containing proteins were present in the insoluble fraction **(Figure S2C).**

**Figure 2:**
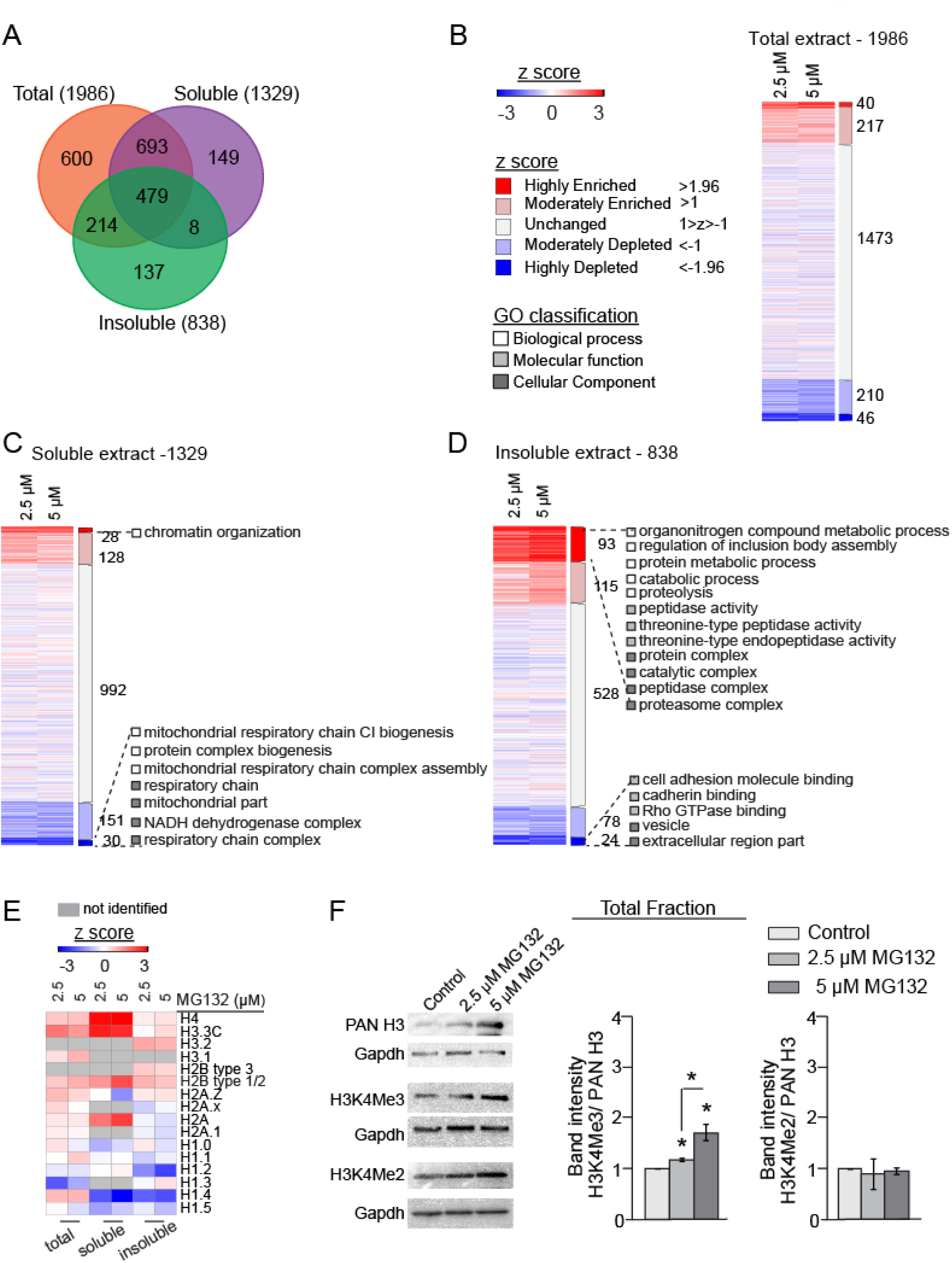
Proteome reorganization in response to proteasome inhibition for 8 hr. (A) Venn diagram showing overlap between the quantified proteins in total, soluble and insoluble fractions of MG132-treated Neuro2a cells. (B) z-score distribution of the quantified proteins in the total fraction. No GO class significantly represented the enriched or depleted proteins. (C and D) z-score distribution of the quantified proteins in the soluble fraction and the insoluble fraction. Significant GO classes representing the highly enriched and highly depleted proteins are shown. Colour scheme similar as **Figure 2B**. (E) Heatmap showing z-score distribution of histones in the proteome fractions. (F) Western blots showing Histone H3, H3K4Me3 and H3K4Me2 levels in the total fraction. Band intensities were normalized to pan Histone H3 protein-levels. Extract of DMSO treated cells served as control. Error bars indicate SDs from at least three independent experiments. ^*^ indicates p<0.05 by Student’s t-test. See also **Figure S2** and **Table S1 and S2**.

Next, we divided the quantified proteins in multiple groups according to their redistribution profile in the MG132 treated cells. A cut-off of 95% confidence, which corresponds to z-scores beyond ±1.96 in either 2.5 or 5 μM treatment, revealed 40 and 28 proteins to be highly enriched and 46 and 30 proteins to be highly depleted in total and soluble fractions, respectively as compared to DMSO control cells **(Figure 2B and 2C; Table S1A and S1B)**. As insoluble fraction showed a skewed distribution of z-scores, it was normalized with mean and standard deviation of total fraction (see methods for detail). Accordingly, we identified 93 (almost 11%) proteins with z-score beyond +1.96 indicating a tendency of many proteins towards insolubility with MG132 treatment. In contrast, 24 proteins showed z-scores below −1.96 **(Figure 2D; Table S1C)**.

Not surprisingly, ubiquitin-load was found to be highly increased in all the proteome fractions of MG132 treated cells. The redistributed proteins in total fraction did not represent any significant Gene Ontology (GO) class (**Figure 2B**). The highly enriched proteins in soluble fraction contained many chromatin associated proteins including Histone H3 and H4. On the other hand, the highly depleted proteins in this fraction included multiple mitochondrial respiratory chain complex (RCC) and mitoribosome subunits (**Figure 2C; Table S1B**).

The insoluble fraction displayed increased abundance of many protein degradation machinery components including proteasome subunits and ubiquitin ligases with MG132 treatment (**Figure 2D**), similar to that described earlier (Wilde et al., 2011). However, only few chaperones (Hsps) were enriched in this fraction suggesting that we have studied a very early time point when even chaperones were not engaged in the refolding of insoluble proteins. The other proteins enriched in insoluble fraction included multiple RCC subunits (detail discussion in following sections) (**Table S1C**). On the other hand, several proteins that are catalogued as ‘exosome components’ or ‘secreted’ in the Exocarta database (Mathivanan and Simpson, 2009) were among the highly depleted proteins in insoluble fraction (**Figure 2D; Table S1C)**. These included α-synuclein (Snca) that is known to be secreted by both vesicle-mediated and unconventional ways (Emmanouilidou et al., 2010; Lee et al., 2016). We confirmed the dose-dependent secretion of overexpressed Snca by proteasome-inhibited cells by analyzing the cell culture media (**Figure S2D**). Other putative secreted proteins that were depleted from insoluble fraction included Rho GTPase binding proteins Arhgdia, Iqgap1, Pfn1 and Flna, cytoskeletal protein Tln1 etc. Similarly, Histone protein Hist1h1e, cytoskeletal protein Marcksl1, small mitoribosomal subunit Mrps6 and ER membrane component Tmed4 are also listed as secreted proteins and their depletion was noted in soluble fraction. Several Exocarta-catalogued secreted proteins were identified as highly depleted in total fraction as well. These are stress signaling protein Map4k4, Rho GDP dissociation inhibitor (GDI) beta (Arhgdib), thiol protease inhibitor (Cstb) and cytoskeleton associated proteins Fn1, Jup and Marcks (**Table S1**). Together, these observations suggest that secretory mechanisms were over-activated in the proteasome-inhibited cells and could be a major route for destabilized protein clearance (Lee et al., 2016).

As described above, significantly increased load of Histone H3.3, H4 and Histone-modifier protein Mortality factor 4-like 2 (Morf4l2) was observed in the soluble proteome. This suggested several possibilities including reduced degradation of these proteins due to proteasome inhibition, dissociation from the chromatin or increase at the transcription-level etc. (**Figure 2E; Table S2**). Transcriptional activation of Morf4l2, Histone H3.3 and H4 could be ruled out as the mRNA contents were unchanged (**Figure S2E**). Morf4l2 is a known proteasome substrate (Larance et al., 2013). Limited degradation of this protein due to proteasome inhibition explained its over-abundance in soluble as well as in total proteome fraction (**Table S1A and S1B**). Similarly, Histone H3.3 and H4 abundance was also increased in total fraction suggesting these proteins also to be proteasome substrates (**Figure 2E**).

Nevertheless, redistribution of Histone and associated proteins across the proteome fractions strongly suggested transcriptional reorganization events. Different covalent modifications at the amino termini of the histones, including acetylation, phosphorylation and methylation have significant influence on gene-specific transcription regulation (Berger, 2002; Karlic et al., 2010). As Histone H3.3 and H4-levels were elevated, we tested the possibility of consequential increase in some of the post-translational modifications on these histones by a series of western blots. While most of the modification levels remained unchanged (**Data not shown**), Histone H3 was found to be significantly trimethylated at lysine 4 position (H3K4Me3) (**Figure 2F and S2F**). Increase in H3K4Me2 (H3-lysine4-di-methylation) was also noticed but this modification turned out to be an apparent consequence of the overall increase of Histone H3 protein-load (**Figure 2F**). Increased load of both Histone H3 and H3K4Me3 was consistent in another potent proteasome-inhibitor lactacystin treated cells confirming this as a proteasome-dysfunction specific response (**Figure S2G**). H3K4Me3-load was also increased in heat-stressed (42°C for 2 hr) cells but the Histone H3 protein-level was unchanged (**Figure S2H**). All these results suggested potential involvement of H3K4Me3 in the active transcription events following proteotoxic-stresses that may include activation of heat shock response. Indeed, increased H3K4Me3-load at the hsp70-promoter region is reported in heat stressed cells (Kim et al., 2011).

### Aggregation of RCC subunits in proteasome-inhibited cells

#### Redistribution pattern - Total fraction

The other group of proteins that displayed prominent redistribution pattern in proteasome-inhibited cells were mitochondrial inner membrane or matrix localized RCC subunits. Among the 282 quantified mitochondrial proteins in total fraction; 9 were highly enriched (Z-scores>1.96) due to MG132 treatment and included 3 respiratory chain complex I (CI) subunits (**Figure 3A; Table S3A**). When we reduced the statistical stringency of data analysis and considered proteins with 1.0< z-scores<1.96 in either 2.5 or 5 μM MG132 treatment as "moderately enriched" proteins; 54 more mitochondrial proteins populated the list amongst which 14 were RCC subunits (6 CI, 3 CIII, 1 CIV and 4 CV) and 5 were mitoribosome components (**Table S3A**). In addition, 33 mitochondrial inner membrane or matrix proteins including Idh3g, Suclg2 and Pdhx that are TCA cycle components; Timm17b, Timm44 and Tomm20 that participate in the import of RCC subunits, etc were among the moderately enriched proteins. Interestingly, three Complex I (CI) subunits (Ndufa5, Ndufa6 and Ndufb4) and 2 mitoribosomal proteins were found to be slightly depleted (−1.0> z-score>-1.96) in total fraction suggesting either secretion or degradation via unknown proteases. Cochaperone Dnajc15 that works as a negative regulator of mitochondrial supercomplex formation (Porras and Bai, 2015) was also found to be depleted (**Table S3A**).

**Figure 3:**
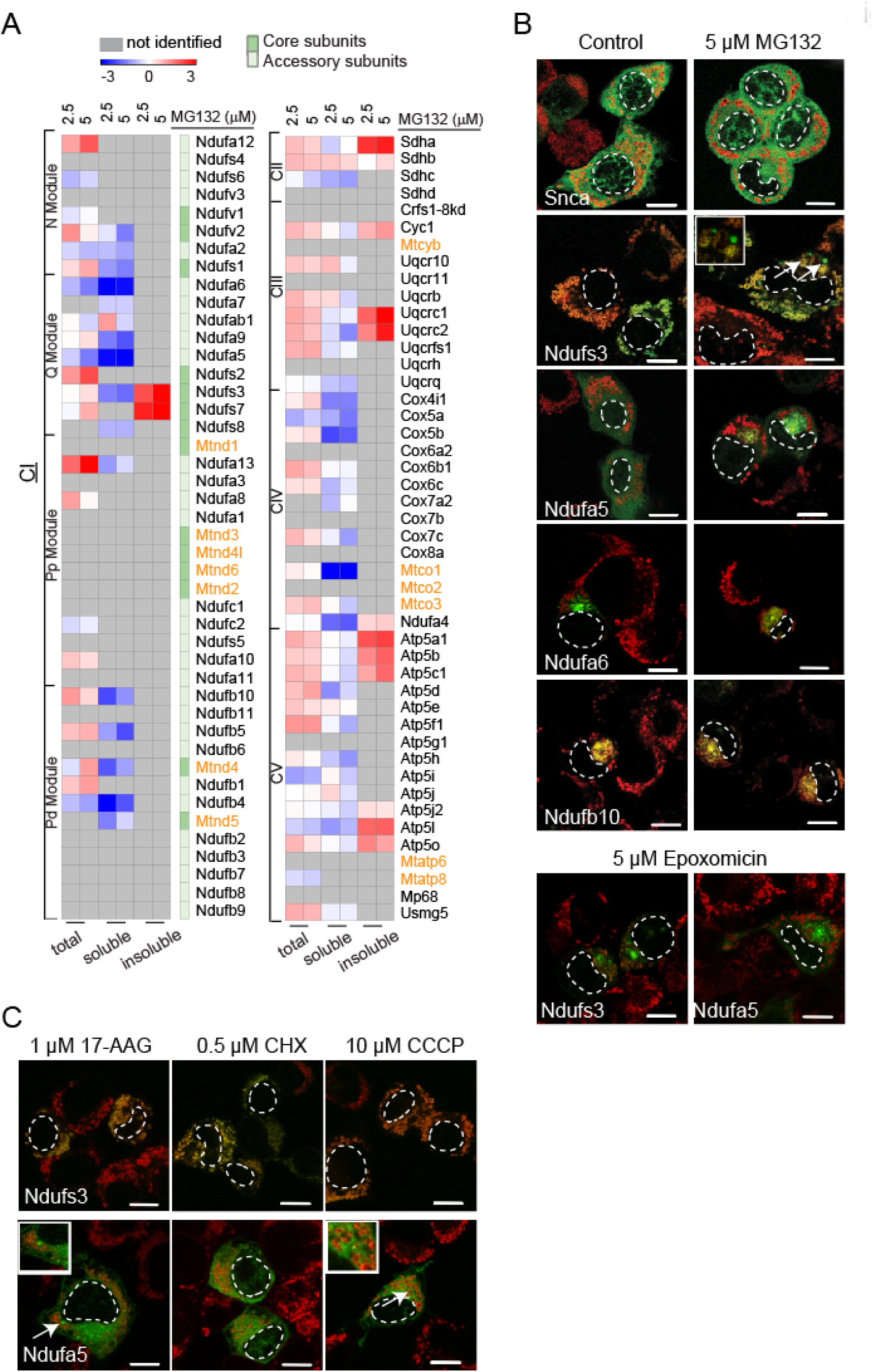
Aggregation of RCC subunits in proteasome-inhibited cells. (A) Heatmap showing z-score distribution of RCC subunits across the proteome fractions of MG132-treated Neuro2a cells for 8 hr. Mitochondria-encoded subunits are shown in orange. (B) Fluorescence micrographs showing IBs formed by overexpressed EGFP tagged RCC subunits with MG132 and epoxomicin. treatment (C) Microscopy images of subunits with Cycloheximide (CHX), 17-AAG and CCCP. Mitochondria are stained by Mitotracker CMXRos (red). Cell fixation was performed using acetone-methanol. Nucleus is shown by dotted lines. Arrows indicate smaller IBs shown in zoomed insets. Control: DMSO treated cell. Scale-bar - 10 μm. See also **Figure S3**, **Table S3, Movies S1 and S2.**

#### Redistribution pattern - Soluble fraction

Among the 187 mitochondrial proteins quantified in soluble fraction, only 1 was found to be highly enriched (z-score>1.96) and 12 moderately enriched (1.0< z-score<1.96) (**Table S3B**). The highly enriched protein was Dnajc19; a putative component of the TIM23-complex-associated import motor that drives transport of RCC precursors into mitochondria (Dennerlein and Rehling, 2015). CI component Ndufab1 was the only slightly enriched RCC subunit in soluble fraction. On the other hand, 15 mitochondrial proteins were highly depleted in the soluble proteome and included 6 CI, 2 CIV and 3 mitoribosome components (**Figure 3A; Table S3B**). TCA cycle component Idh3g was also highly depleted. Fifteen more RCC subunits, 3 mitoribosome components and the supercomplex assembly inhibitor Dnajc15 were among the moderately depleted proteins (**Table S3B**). In absence of transcriptional deregulation as confirmed by RT-PCR experiments (**Figure S3A**), these proteins are either partitioned into the insoluble fraction or secreted or degraded by some unconventional mechanism.

#### Redistribution pattern - Insoluble fraction

In the insoluble fraction, 18 mitochondrial proteins were highly enriched and included 2 CI, 1 CII, 2 CIII and 1 CV subunits. Moderately enriched insoluble proteins contained 26 mitochondrial proteins among which 1 CIII and 4 CV components were present (**Figure 3A**; **Table S3C**). TCA cycle protein Suclg2 was also found to be highly enriched (**Table S3C**). Mitoribosomes localize in close proximity of RCC complexes on the mitochondrial inner membrane and synthesize the 13 core RCC subunits (7 CI, 1 CIII, 3 CIV and 2 CV). As discussed above, we observed multiple mitoribosome components to be increased in abundance in total fraction and decreased in soluble fraction (**Table S3**). Being highly hydrophobic in nature, the core RCC subunits synthesized by mitoribosomes are difficult to identify by mass spectrometry (Miwa et al., 2014) and most of them were not quantified in our experiments. However, when identified, these were also increased in total and depleted in soluble fraction (**Figure 3A**).

Thus, mass spectrometry experiments revealed a distinctive reorganization pattern of mitochondrial RCC subunits as an early event in proteasome-inhibited cells and motivated us to further investigate the nature, extent, and mechanism of the reorganization of RCC subunits and its impact on cellular homeostasis. Fully assembled CI is the largest respiratory complex, composed of 44 subunits and divided in 4 modules. The matrix arm comprises of N and Q modules responsible for NADH dehydrogenase and ubiquinone reductase activity, respectively. Proximal and distal proton pump modules (P_P_ and P_D_) are present in the membrane arm. Except the 7 mitochondrially synthesized subunits, CI components are encoded by nuclear genes, synthesized in the cytoplasm, and imported into the mitochondria in an unfolded state by an ATP-dependent chaperone mediated process (Guerrero-Castillo et al., 2017; Harbauer et al., 2014). As described above, these nuclear-encoded proteins are enriched in total fraction, depleted in soluble fraction and enriched in insoluble fraction (**Figure 3A**) suggesting aggregation of these proteins. To validate, we cloned CI subunit Ndufs3 with EGFP tag in mammalian expression vector. As reported earlier, overexpression of this N-module subunit does not perturb CI assembly and function (Vogel et al., 2007) and did not affect cell viability in our experiments as measured by MTT assay (**data not shown**). After transfection, Ndufs3-EGFP was found to be largely colocalized with mitochondria. Consistent with our mass spectrometry observations, multiple distinct mitotracker negative IBs appeared after treating the cells with MG132 for 8 hr (**Figure 3B; Movie S1**). Also, we observed collapse of mitochondrial morphology along with Ndufs3 aggregates in several MG132 treated cells (**Figure S3B**).

Redistribution pattern of two Q-module subunits Ndufa5 and Ndufa6 was atypical as these were depleted from both total and soluble fractions whereas not identified in the insoluble fraction (**Figure 3A**). When tagged with EGFP and overexpressed in Neuro2a cells, Ndufa5 displayed partial colocalization with mitochondria and otherwise diffused distribution throughout the cells including faint staining in the nucleus. However, upon MG132 treatment it was found to be sequestered in the cytosol nearby nucleus with faint mitotracker staining in the surrounding (**Figure 3B; Movie S1**). EGFP-tagged versions of both Ndufs3 and Ndufa5 were found to be precipitated into the insoluble fraction of MG132 treated cells and ubiquitinated suggesting these proteins to be substrates of the ubiquitin-proteasome system (**Figure S3C and S3D**). Ndufa6 and a P_D_ module subunit Ndufb10 immediately sequestered in the cytoplasm upon transfection and disorganized mitotracker staining suggested collapsed mitochondrial structure in the vicinity (**Figure 3B; Movie S1**).

At high dose, MG132 blocks matrix localized Lon protease in isolated mitochondria that may contribute to the destabilization of the RCC subunits. Nevertheless, Ndufs3 and Ndufa5 were found to form aggregates in presence of epoxomicin (**Figure 3B**), a proteasome inhibitor without any impact on Lon Protease (Bayot et al., 2008; Bezawork-Geleta et al., 2015), ruling out that possibility. Also, a non-mitochondrial protein α-synuclein was distributed diffusedly in the MG132 treated cells without affecting mitochondrial morphology suggesting that proteasome inhibition specifically destabilizes the nuclear-encoded RCC subunits (**Figure 3B; Movie S1**). Perturbation of other arms of proteostasis network using translation elongation blocker cycloheximide, Hsp90 inhibitor 17-AAG, and mitochondrial protein translocation blocker CCCP did not result in similar massive aggregation of Ndufs3 and Ndufa5. However, in case of Ndufa5, few smaller aggregate-like structures were observed in 17-AAG and CCCP treated cells (**Figure 3C; Movie S2**) that are morphologically very similar to phase-separated droplets of Tau (Vidoni et al., 2017). These observations suggested that RCC subunits are aggregation-prone in presence of multiple proteotoxic stresses. As the local concentration of these proteins is significantly increased due to reduced degradation in proteasome-inhibited cells, they collapse into large aggregates.

### Role of LCR containing N-terminal regions of RCC subunits in aggregation

Being components of large and abundant multi-subunit complexes, RCC subunits are likely to contain aggregation-prone interaction surfaces (Pechmann et al., 2009). Hence, we investigated whether the insolubility of these vulnerable subunits is determined by common physicochemical signatures or are they of unrelated sequence and structures and randomly aggregated. High iBAQ values suggested that these are abundant proteins. These are also physicochemically distinct with low molecular weight and basic pI (**Figure S4A**). Out of the 21 nuclear-encoded RCC subunits identified to be depleted in soluble fraction, 14 contained arginine-rich N-terminal predicted mitochondrial targeting signal (MTS) that majorly contribute for the basic pI of these proteins (**Figure 4A, 4B and S4A**). In addition, the MTS sequences were overrepresented by alanine, leucine and serine residues (**Figure 4B**). Also, 7 of these MTS sequences were predicted to contain low complexity regions (LCR) (**Figure 4A**). For the remaining 7 RCC subunits that are depleted in soluble fraction but without signal sequences, we inspected the first 30 amino acids from the N-terminal which is the average length of MTS. These sequences were not rich in arginine but few of them contained LCRs and had higher percentage of threonine and serine residues that are potential targets for phosphorylation (**Figure 4A, 4B and S4B**). Interestingly, phosphorylation within the LCR has been identified as triggering-factor for aggregation of Tau (Ambadipudi et al., 2017).

**Figure 4:**
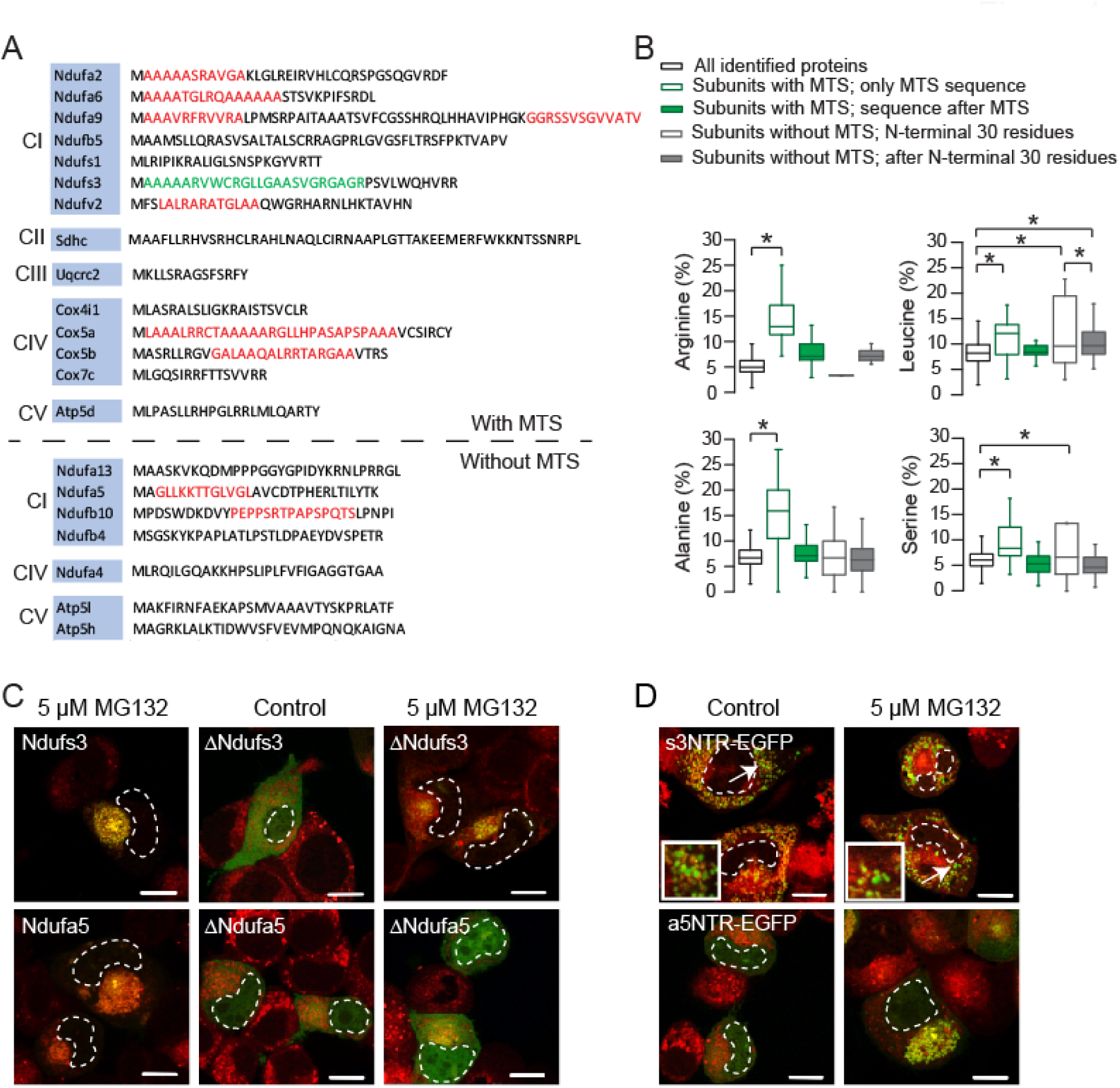
Role of LCR containing N-terminal regions of RCC subunits in aggregation. (A) Low complexity regions (LCRs) in the predicted MTS or N-terminal 30 residues of RCC subunits depleted in the soluble fraction. High confidence LCRs are highlighted in red; low confidence in green. (B) Box plots showing amino acid composition of MTS of RCC subunits. MTS or N-terminal 30 residues and rest of the protein-sequences are grouped separately. ^*^ indicates p<0.05 by one-way ANOVA, Bonferroni’s post hoc test. (C) Fluorescence micrographs of EGFP tagged RCC subunits without N-terminal 30 amino acid sequence. (D) Fluorescence micrographs of EGFP with N-terminus sequence of RCC subunits. Mitochondria are stained by Mitotracker CMXRos (red). Control: DMSO treated cells. Cell fixation was performed using paraformaldehyde. Nucleus is shown by dotted lines. Arrows indicate smaller IBs shown in zoomed insets. Scale-bar - 10 μm. See also **Figure S4** and **Movie S3**.

In addition to the depleted proteins in soluble fraction, N-terminal MTS sequences of many of the nuclear-encoded RCC subunits enriched in total and insoluble fractions also contained LCRs (**Figure S4C and S4D**). Despite some random fluctuation events, the N-terminal amino acid stretches of Ndufs3, Ndufa5 and Ndufa6 displayed reduction of the radius of gyration (R_g_) values by 50 ns in molecular dynamics simulation (**Figure S4E**) suggesting conformational compactness of these peptides with time (Amin et al., 2014; Clark, 2005; Ghattas et al., 2017). N-terminus of Ndufs3 contained a weakly predicted LCR but a strong MTS. Mitochondrial localization of this protein was lost upon removal of N-terminus and ∆Ndufs3 was diffusedly distributed throughout cytoplasm but formed aggregates upon MG132 treatment (**Figure 4C; Movie S3**). When we deleted the LCR containing N-terminal 30 amino acids from Ndufa5 (∆Ndufa5), the truncated version did not form large aggregates in MG132 treated cells (**Figure 4C**), rather few small aggregate-like structures were still present at different z-planes (**Movie S3)**. On the other hand, addition of MTS of Ndufs3 at the N-terminus targeted EGFP (s3NTR-EGFP) to mitochondria along with some aggregate-like puncta not co-localising with mitotracker. Moreover, N-terminus of Ndufa5 was sufficient to trigger massive aggregation of EGFP (a5NTR-EGFP) upon MG132 treatment (**Figure 4D; Movie S3**). Thus, our data suggest that LCRs in the RCC subunits are capable to trigger their aggregation, however, contribution from other sequence features cannot be ruled out. Contextually, liquid to liquid and liquid to solid phase separation of LCR containing proteins has recently been identified as an adaptive cellular mechanism in proteotoxic conditions (Franzmann et al., 2018; Mitrea and Kriwacki, 2016; Molliex et al., 2015).

### Consequence of the aggregation of RCC subunits - mitochondrial morphology and function

In addition to the RCC subunits, many other nuclear-encoded mitochondrial proteins are expected to be redistributed in MG132 treated cells as they may contain LCR containing MTSs for translocation. Indeed, the z-score distribution pattern showed increased abundance of many mitochondrial proteins in total fraction. Moreover, many of them were depleted in the soluble fraction whereas increased in insoluble fraction (**Figure 5A; Table S3**). In comparison, z-score profiles of other organelle-components like nucleus or endoplasmic reticulum did not display any pattern (**Figure 5A**). Thus, the redistribution profile due to MG132 treatment was mitochondria-specific and suggested that many mitochondrial precursor proteins could be potential proteasome substrates; RCC subunits being the most prominent examples.

**Figure 5:**
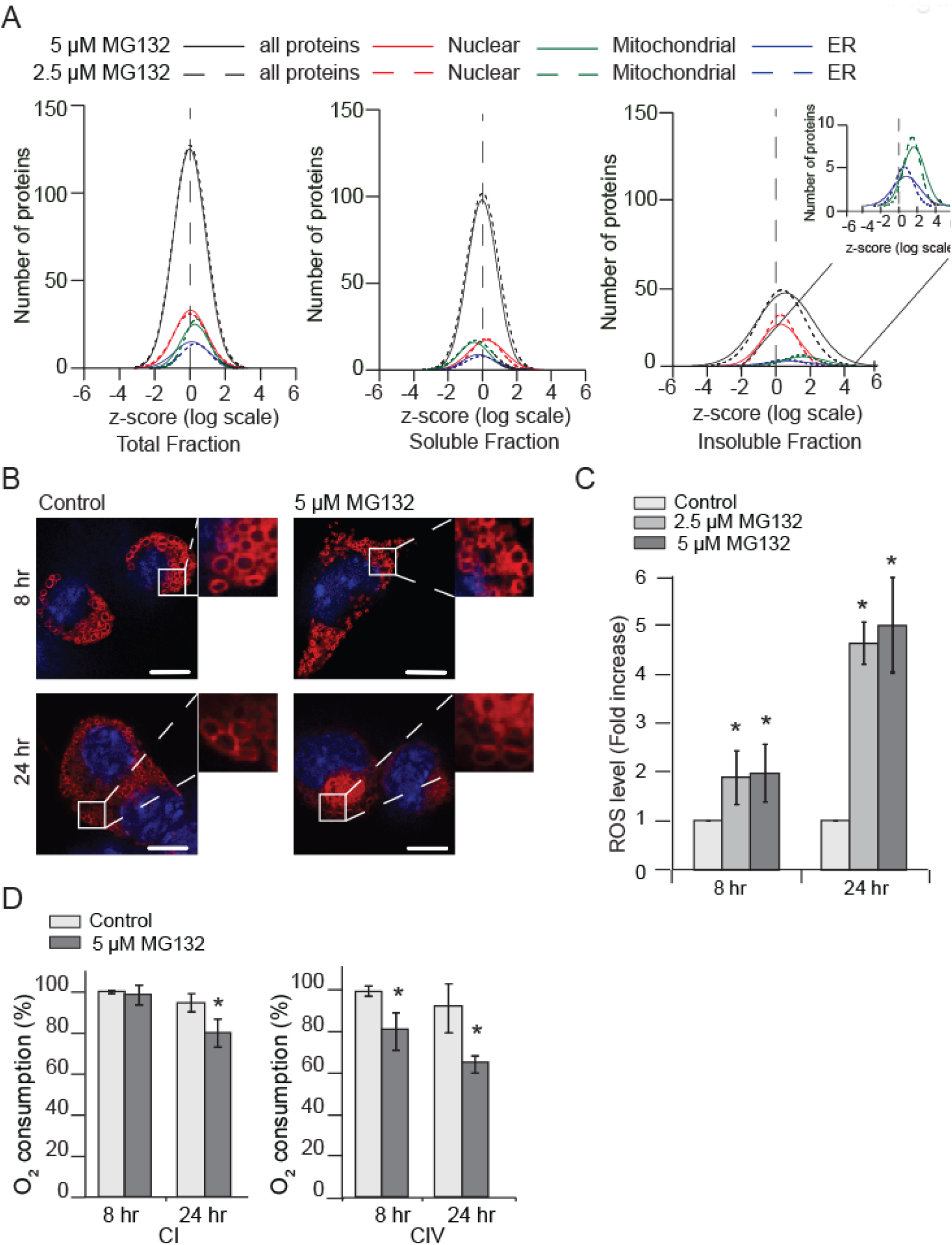
Consequence of aggregation of RCC subunits - mitochondrial morphology and function. (A) Histogram distribution of z-scores of organelle-proteins (as curated from Uniprot) with MG132 treatment across the proteome fractions. Rightmost inset shows zoomed z-score distribution for organelle-proteins in the insoluble fraction. (B) A closer look at mitochondrial morphology in untransfected MG312-treated cells for different time-points. Mitochondria are stained by Mitotracker CMXRos (red). Cell fixation was performed using acetone-methanol. Control: DMSO treated cells. Scale-bar - 10 μm. Digitally zoomed sections are shown as insets. (C) ROS production by MG312-treated cells for indicated time-points. (D) Complex I and IV activity was determined on digitonin-permeabilized cells in mitochondrial respiration medium Mir05 using Oxygraph-2K. Control: DMSO treated cells. Error bars indicate SDs from at least three independent experiments. ^*^ indicates p<0.05 by Student’s t-test. See also **Figure S5**.

Strikingly, mitochondrial structure and function was almost unperturbed despite of this extensive proteome disorganization. No change in mitochondrial morphology was observed, only mild membrane depolarization was detected and ROS level was increased only 2-fold by 8 hr of MG132 treatment (**Figure 5B and 5C**). Mitochondrial unfolded protein response (UPRmt) was slightly elevated as evident by moderate increase of Hsp60; an Atf 5-dependent mitochondrial chaperone (Fiorese et al., 2016), in insoluble fraction but not in total or soluble fraction (**Table S1**). Nevertheless, mitochondrial morphology was completely collapsed when we prolonged the treatment for 24 hr and severely depolarization was noted with 5-fold increase in ROS levels (**Figure 5B and 5C**). To probe further, we performed high-resolution respirometry in permeabilized Neuro2a cells to estimate the individual contribution by respiratory chain complexes in cellular respiration. Although many CI subunits were found to be redistributed, CI dependent electron transfer was not significantly affected by 8 hr of MG132 treatment as evident by no change in oxygen flux. CII function was also not perturbed but significant drop (~20%) was found for oxygen consumption by CIV. CI activity was indeed compromised with prolonged proteasome inhibition as evident by a moderate and statistically significant decrease in oxygen consumption after 24 hr (**Figure 5D and S5**).

### Consequence of the aggregation of RCC subunits - destabilization of sub-assemblies of respiratory complexes

Unperturbed CI activity suggested presence of functional respiratory complex at the mitochondrial inner membrane at the early stage of proteasome inhibition. Therefore, we speculated some deregulations in the assembly process due to aggregation of the subunits before their incorporation into the functional respiratory complexes. Recent complexome profiling experiments revealed that multiple small intermediates follow a stepwise mechanism to assemble into the four modules of CI (Guerrero-Castillo et al., 2017). These small intermediates are mainly composed of nuclear encoded subunits to which mitochondrially coded subunits associate to form larger or fully assembled complex. Interestingly, most of the aggregation-prone CI subunits (including Ndufs3, Ndufa5, Ndufb10 etc.) were found to be components of different small assembly intermediates that are translated in the cytosol suggesting a perturbation in the CI-assembly process by 8 hr of proteasome-inhibition (**Figure 3A**). Hence, we analyzed digitonin-solubilised mitochondrial fractions on a Blue Native-PAGE (BN-PAGE) that separates the mitochondrial protein complexes according to molecular mass. The gel was divided in 20 different slices and quantitative mass spectrometry was performed to investigate the distribution of CI subunits across the assembly intermediates (**Figure S6A**). Slice 4-7, the region that corresponds to ~ 800 - 1000 kDa, contained almost all the subunits of CI in high abundance indicating the fully assembled CI (**Figure 6A; Table S4**). Slice 9-15 contained multiple Q and P_D_ module components suggesting presence of intermediate assembly products. All the core subunits of N module (Ndufv1, Ndufv2 and Ndufs1) were found in slice 16-20 (~ 150-250 kDa) along with the Q and P_D_ module components suggesting presence of multiple small assembly units (**Figure 6A; Table S4A**). SILAC based quantitation indicated significant reduction of these small assembly intermediates (slice 16-20) in MG132 treated cells by 8 hr in dose-independent manner (**Figure 6A and S6B**). However, the fully assembled CI (Slice 4-7) was still intact by the short-term treatment.

**Figure 6:**
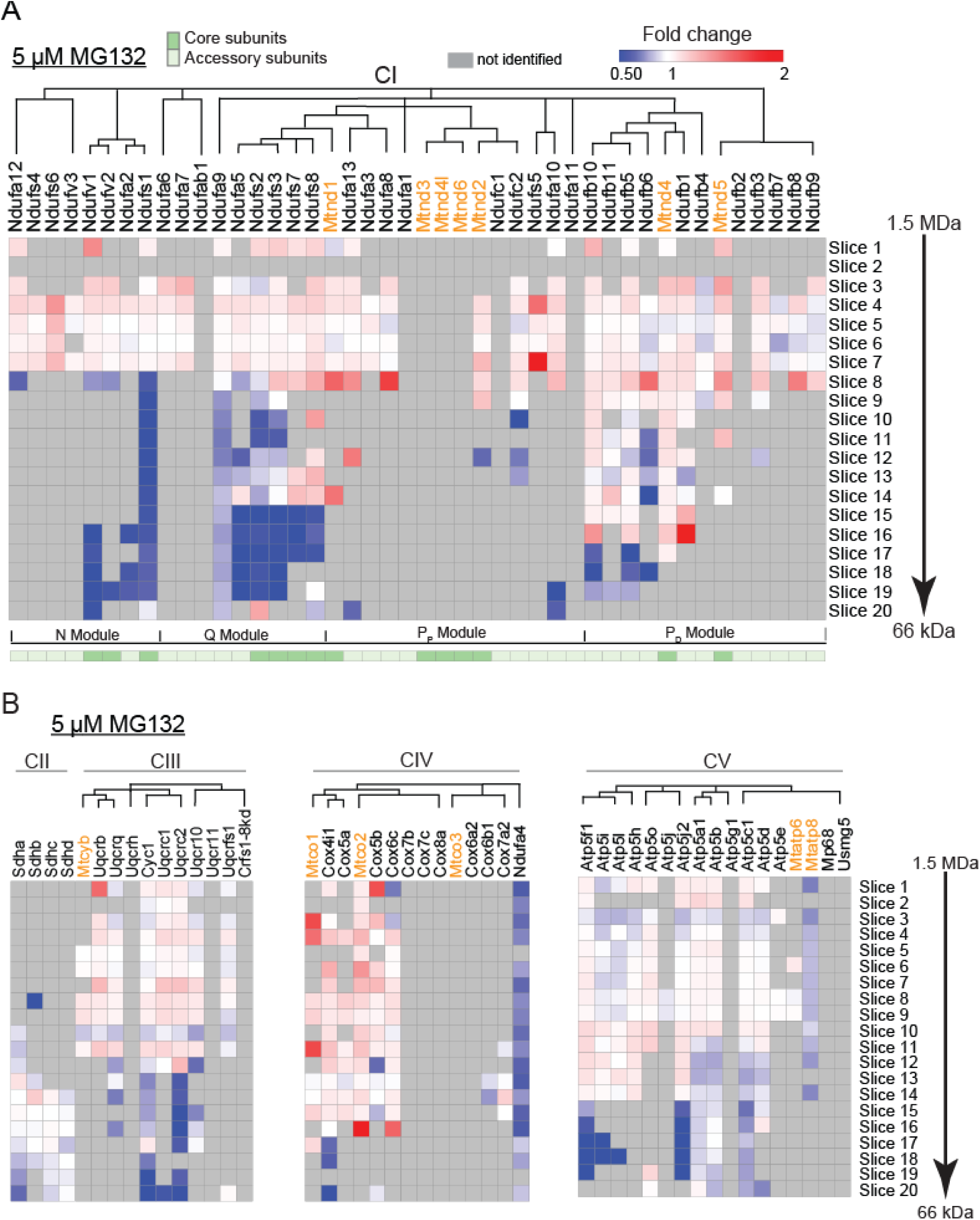
Consequence of aggregation of RCC subunits - destabilization of sub-assemblies of respiratory complexes. (A) Alteration of abundance of CI subunits at different slices of BN-PAGE in MG132-treated cells (8 hr) as determined by SILAC-based fold changes over control DMSO treated cells. The subunits are clustered as per CI assembly pathway described by Guerrero-Castillo et al. (Guerrero-Castillo et al., 2017). Mitochondria-encoded subunits are shown in orange. (B) Alteration of abundance of CII, CIII, CIV and CV subunits at different slices of BN-PAGE in MG132-treated cells (8 hr) as determined by SILAC-based fold changes over control DMSO treated cells. The subunits are clustered as per the assembly pathways described previously (Vidoni et al., 2017, Fujikawa et al., 2015, Fernandez-Vizarra et al., 2015). Mitochondria-encoded subunits are shown in orange. See also **Figure S6 and Table S4**.

Unlike CI, other respiratory complexes are constituted by less number of subunits and their assembly mechanisms have not been investigated that extensively. All 4 CII subunits were abundant in slice 14-17 (~ 150 - 500 kDa) and unaffected by short-term MG132 treatment (**Figure 6B; Table S4B**). CIII is composed of one mitochondrial and ten nuclear-encoded subunits and was majorly found in slice 4-6 (~ 900 kDa) and slice 8-11 (~ 500-700 kDa). Nuclear-encoded CIII subunits Cyc1, Uqcrc1 and Uqcrc2 are known to constitute a sub-complex that binds to membrane bound Mt-Cyb at an intermediary step of assembly process (Fernandez-Vizarra and Zeviani, 2015). All the components of this sub-complex were identified together in slice 19-20 of the BN-PAGE and were depleted in the proteasome inhibited cells by 8 hr (**Figure 6B**). To note here, both Uqcrc2 and Uqcrc1were found to be decreased in soluble and increased in abundance in insoluble fraction of the MG132 treated cells (**Figure 3A**).

CIV subunits showed a wide distribution in the BN-PAGE; in slice 4 (~ 1000 kDa), 8-9 (~ 700 kDa), and 13-15 (~ 480 kDa) suggesting its possible association with other respiratory complexes towards the formation of supercomplexes (**Figure 6B**; **Table S4B**). Assembly of CIV begins with an intermediate containing Cox4i1 and Cox5a (Vidoni et al., 2017) and both these proteins were found to be depleted in soluble fraction (**Figure 3A**). We could not identify Cox5a in the BN PAGE analysis; however, protein level of Cox4i1 was significantly reduced in slice 20 suggesting a disruption of this intermediate by 8 hr of MG132 treatment (**Figure 6B**). Another protein Ndufa4, that has been recently identified to be critical for both structural and functional stability for CIV (Balsa et al., 2012), was found to be moderately reduced throughout the BN PAGE and highly reduced in soluble fraction of proteasome-inhibited cells (**Figure 6B**). Thus, drop in oxygen consumption by CIV by short-term MG132 treatment could be attributed to unavailability of functional Ndufa4.

CV subunits were most abundant in slice 8-9 (~ 700 kDa) that corresponds to the holocomplex (**Figure 6B**; **Table S4 Table S4B**). An assembly intermediate for the stator stalk of CV containing Atp5f1, Atp5l and Atp5i has been reported recently (Fujikawa et al., 2015). We observed reduction in the abundance of all these subunits at ~ 150-250 kDa of BN PAGE (Slice 16-18) indicating loss of this assembly unit with short-term proteasome inhibition (**Figure 6B**). The combined mass of these 3 subunits is ~ 44 kDa. Hence, its appearance at ~ 150-250 kDa region suggested presence of other CV components as constituent of the same assembly intermediate. We identified Atp5j2 and Atp5c1 in the same region and abundance of both these components were also reduced with MG132 treatment (**Figure 6B**). Strikingly, Atp5f1, Atp5l and Atp5i were also found to be redistributed across the proteome fractions (**Figure 3A**).

## DISCUSSION

The dynamics of the cellular proteome is slowly damaged by continuous accumulation of misfolded insoluble proteins due to perturbation of various nodes of proteostasis network. At the onset, cells attempt to trigger defence mechanisms but in long-term lose the capacity to react and succumb. Snapshots of early proteome destabilization events provide an understanding of the seeding deleterious events that lead to the collapse of cellular processes. Our study revealed precipitation of nuclear-encoded RCC subunits as one of the earliest proteotoxic signatures in proteasome-inhibited cells when even activation of autophagy was not prominent rather secretion was more active mechanism for clearance of destabilized proteins.

Understanding of structure, function or assembly of RCC subunits is a subject of intense research. It appears to be a multistep process and RCC components are not active individually but only when assembled. The individual respiratory complexes further assemble into supercomplexes or respirasomes on the tip of the cristae in the mitochondrial inner membrane offering several structural or functional advantages including prevention of destabilization of RCC components, enhancement of electron transport efficiency, substrate channelling etc. (Gu et al., 2016; Guo et al., 2016; Stroud and Ryan, 2013). The assembly process begins with association of multiple matrix arm subunits in small intermediates acting as scaffolds for the coordinated and sequential incorporation of other partially assembled subunits (Melber and Winge, 2016). Most of these nuclear-encoded small assembly intermediates include at least one subunit with MTS at the N-terminus consisting of LCRs over-representing arginine, alanine, leucine or serine residues. Increased number of positively charged amino acids at the N-terminus could be a requirement for interaction with the translocon systems on the mitochondrial membranes or with the lipids like cardiolipin (Harbauer et al., 2014; Pfeiffer et al., 2003); however, such stretches are known to be aggregation prone (Boeynaems et al., 2017). Poly-alanine containing segments have also been shown to accelerate protein aggregation (Pelassa et al., 2014). Moreover, these RCC subunits are known as supersaturated proteins because of their high abundance and intrinsic insolubility (Ciryam et al., 2013). These mitochondrial proteins clumped together into aggregates both inside and outside mitochondria in the proteasome-inhibited cells as the local concentration was further amplified but chaperone-capacity was not sufficient. The LCRs at the N-terminus of these proteins could be potentially involved in seeding this phase separation but contribution from the rest of the protein sequences may also be required for the conformational collapse. As per complexome profiling experiments, the pre-existing fully assembled respiratory complexes were unhurt by the aggregation events keeping the respiratory activities almost unperturbed. Thus, our data indicate only limited threat to mitochondrial homeostasis at short-term but predict potential damages in the association of RCC subunits into the fully assembled complexes during prolonged proteasome inhibition (**Figure 7**).

**Figure 7:**
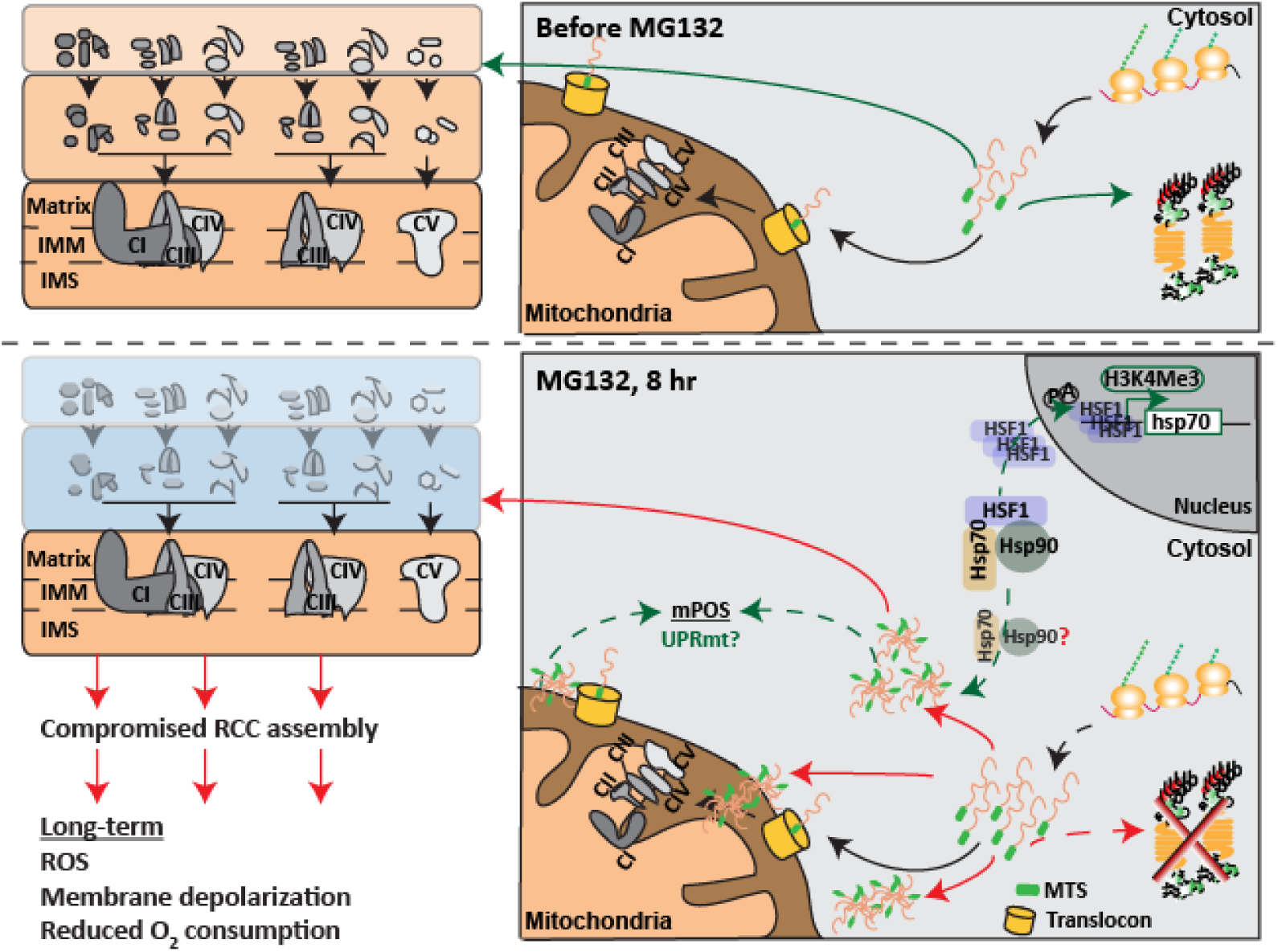
Model of aggregation of RCC subunits and destabilization of respiratory complex assembly in proteasome-inhibited cells. In absence of proteasome inhibition, nuclear encoded RCC subunits are translocated to the mitochondria and assembled into respiratory complexes after cleavage of the MTS. Upon proteasome inhibition, the aggregation-prone RCC subunits accumulate in the cell and form IBs. This leads to disappearance of small and intermediate assembly units of the respiratory complexes that affects mitochondrial respiratory activities in long-term. We propose that these destabilized RCC subunits participate in Hsf1-mediated transactivation of heat shock proteins by sequestering cytosolic Hsp70 into the insoluble fraction.

Progressive failure of both mitochondrial respiration and protein degradation is known since long time in many age-related disease models; however, the molecular link was not adequately understood (Ross et al., 2015). Our work highlights that connection, indicates that cytosolic turnover of multiple RCC subunits is dependent on the ubiquitin-proteasome system and reveals that a collapse in the dynamics of the RCC assemblies plays as catalyst in the gradual loss of respiratory activities and excessive ROS production in the proteasome-inhibited cells (**Figure 7**). Accumulation of ROS slowly generates oxidative stress and affects folding of other proteins including proteasome subunits etc. and begins to damage multiple cellular functions (Ross et al., 2015). In addition, reduced respiratory functions in these cells could chronically impair the ATP-dependent chaperone mediated translocation of unfolded precursors of nuclear-encoded RCC subunits to mitochondria (Guerrero-Castillo et al., 2017). Hence, understanding the dynamics, rate-limiting steps, and identifying the specific chaperones and ubiquitin ligases involved in the transport, assembly and turnover of the RCC subunits might provide valuable insights on the interconnectedness between proteostasis and bioenergetics and their relative contributions in aging or age-related diseases. In this context, reorganization of RCC subunits in other proteotoxic stress conditions including heat stress, amyloid toxicity etc. will be interesting to investigate.

Recent evidences also suggest that mitochondrial-precursor proteins accumulate in the cytosol or mitochondria of yeast cells upon proteasome inhibition triggering a threat to respiratory functions, a phenomenon famed as mitochondrial precursor over-accumulation stress (mPOS) (Wang and Chen, 2015). This triggers activation of multiple adaptive pathways including mitochondrial unfolded protein response (UPRmt), unfolded protein response activated by mistargeting of mitochondrial proteins (UPRam) etc. While UPRmt stabilizes the proteome by elevating mitochondrial chaperone and protease levels; UPRam attenuates protein-synthesis and triggers proteasome-mediated degradation of mistargeted mitochondrial precursors (Haynes and Ron, 2010; Wrobel et al., 2015). We did not observe substantial elevation of UPRmt proteins in our proteomics experiments. Rather, our data indicate that aggregation-prone RCC subunits Ndufa5 and Ndufa6 may be cleared from the MG132 treated cells by unconventional secretion mechanisms (Durieux et al., 2011; Lee et al., 2016). Also, we found that destabilization of the RCC subunits happened together with transcriptional activation of Hsf1-dependent cytosolic stress response. Readjustment of Histone landscape is supportive of this active transcription. In order to re-establish proteostasis, these insoluble RCC subunits may be directly involved in HSR transactivation by titrating the chaperones associated with Hsf1 in the cytosol but remains to be investigated in detail (**Figure S7**).

## SUPPLEMENTARY FIGURES

**Figure S1:**
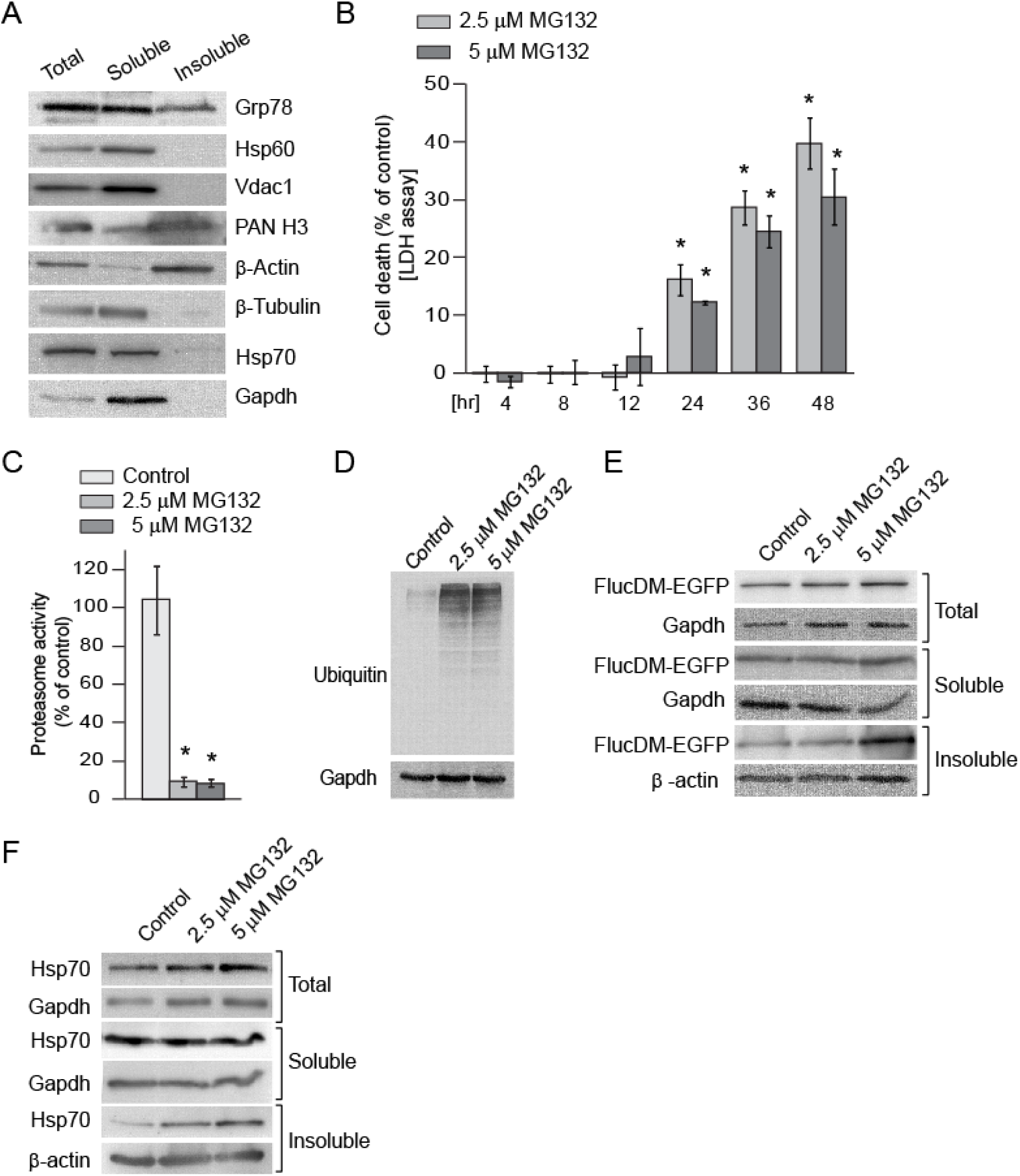
Destabilization of chaperone-dependent proteins in Neuro2a cells by short-term proteasome-inhibition. Related to Figure 1. (A) Western blots displaying the abundance of marker proteins of different cell-organelles in total, soluble and insoluble fraction of Neuro2a cells. (B) Cell toxicity assay. Neuro2a cells were incubated with MG132 for different time durations and cell death was estimated by LDH assay. Control: DMSO treated cells. Error bars indicate SDs from at least three independent experiments. ^*^ indicates p<0.05 by Student’s t-test. (C) Proteasome activity. Neuro2a cells were treated with indicated concentrations of MG132 for 8 hr and proteasome activity was measured by the protocol described in methods. Control: DMSO treated cells. Error bars indicate SDs from at least three independent experiments. ^*^ indicates p<0.05 by Student’s t-test. (D) Ubiquitinated protein-load. Neuro2a cells were treated as described in (C); soluble extract was prepared and western blotting was performed using anti-ubiquitin and anti-GAPDH. (E) Abundance of overexpressed FlucDM-EGFP in total, soluble and insoluble fraction with 8 hr MG132 treatment. (F) Abundance of HSP70 in total, soluble and insoluble fraction of MG132 treated FlucDM-EGFP over-expressing Neuro2a cells (8hr). Gapdh served as loading control for the total and soluble fraction; β-actin for the insoluble fraction.

**Figure S2:**
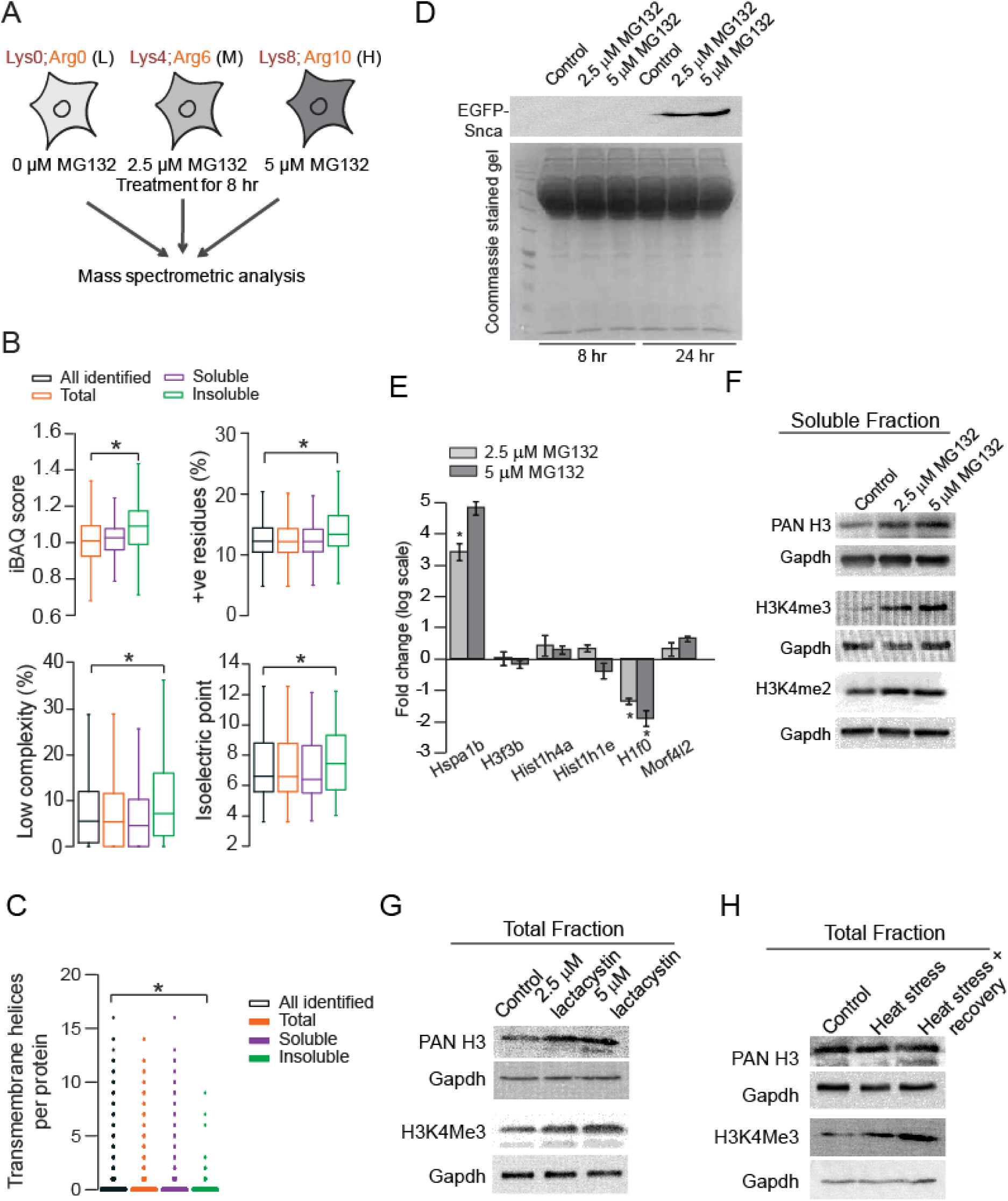
Proteome reorganization in response to proteasome inhibition. Related to Figure 2. (A) Schematic describing the SILAC-based quantitative mass spectrometry strategy to analyze the total, soluble and insoluble fraction of MG132 treated Neuro2a cells. (B) Box plots showing physicochemical properties of the proteins identified in total, soluble and insoluble fraction. iBAQ indicates intrinsic abundance of proteins as calculated from mass spectrometry data from the control DMSO treated cells. ^*^ indicates p<0.05 by one-way ANOVA, Bonferroni’s post hoc test. (C) Box plots showing predicted transmembrane helix content of the proteins identified in total, soluble and insoluble fraction. ^*^ indicates p<0.05 by one-way ANOVA, Bonferroni’s post hoc test. (D) Secretion of EGFP-tagged Snca from MG132 treated cells. Cells were transfected with EGFP-tagged Snca and treated with MG132 for 8 and 24 hr. Culture medium was loaded on SDS-PAGE and western blot was performed with anti-EGFP. A coommassie stained gel loaded with same volume of culture media served as loading control. (E) Fold changes of mRNA-levels of candidate histone genes as determined by real-time PCR. hspa1b (Hsp70) and morf4l2 represented stress induced gene and histone-associated gene, respectively. h3f3b - Histone H3, hist1h4a - Histone H4, hist1h1e - Histone H1.4, h1f0-Histone H1.0. Fold change was calculated against mRNA levels in DMSO treated cells. Error bars indicate SDs from at least three independent experiments. ^*^ indicates p<0.05 by Student’s t-test. (F) Western blots showing Histone H3, H3K4Me3 and H3K4Me2 levels in soluble fraction of MG132 treated Neuro2a cells (8 hr). (G) and (H) Immunoblots showing Histone H3 and H3K4Me3 levels in total fraction of lactacystin treated (8 hr) and heat-stressed (42°C for 2 hr) Neuro2a cells. Control: DMSO treated cells Gapdh level served as loading control

**Figure S3:**
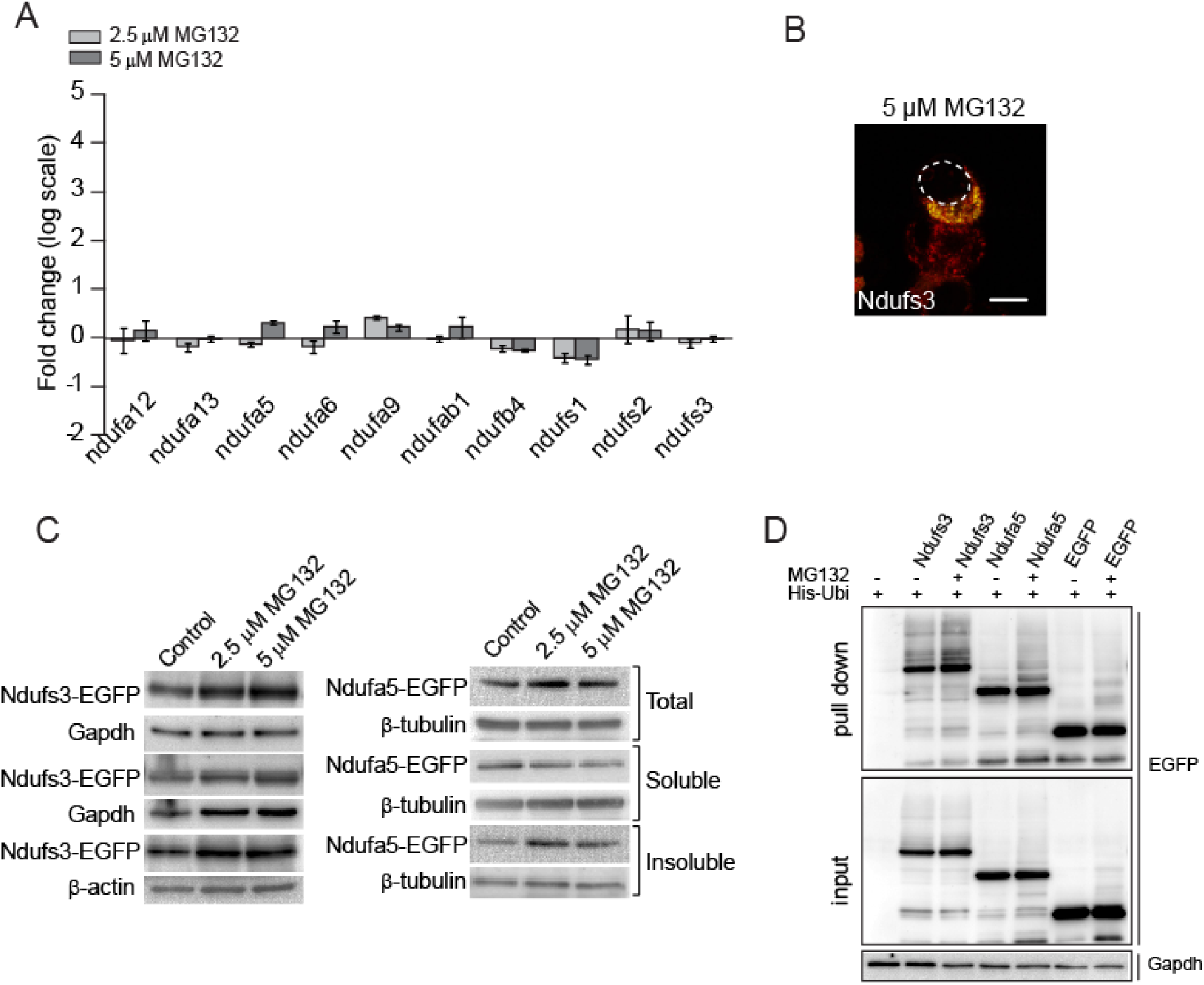
RCC subunits in MG132 treated Neuro2a cells for 8 hr. Related to Figure 3. (A) Fold change of mRNA-levels of candidate RCC subunits with MG132 treatment as determined by real-time PCR. Data were normalized to Gapdh mRNA levels and fold change calculated against DMSO control cells. Error bars indicate SDs from at least three independent experiments. (B) Fluorescence micrographs showing large IBs formed by overexpressed Ndufs3-EGFP. (C) Redistribution of overexpressed Ndufs3-EGFP and Ndufa5-EGFP in total, soluble and insoluble fraction. Gapdh served as loading control for total and soluble fraction; β-actin for insoluble fraction for Ndufs3-EGFP while β-tubulin for Ndufa5-EGFP. (D) Ubiquitylation of RCC subunits. Neuro2a cells were transfected with His-ubiquitin (His-Ubi) and Ndufs3-EGFP or Ndufa5-EGFP or EGFP (as indicated in the figure). After 24 hr cells were exposed to 5 μM MG132 or DMSO. Cells were lysed and His-Ubi pull-down performed. Samples were separated by SDS-PAGE and immunoblotted with anti-EGFP.

**Figure S4:**
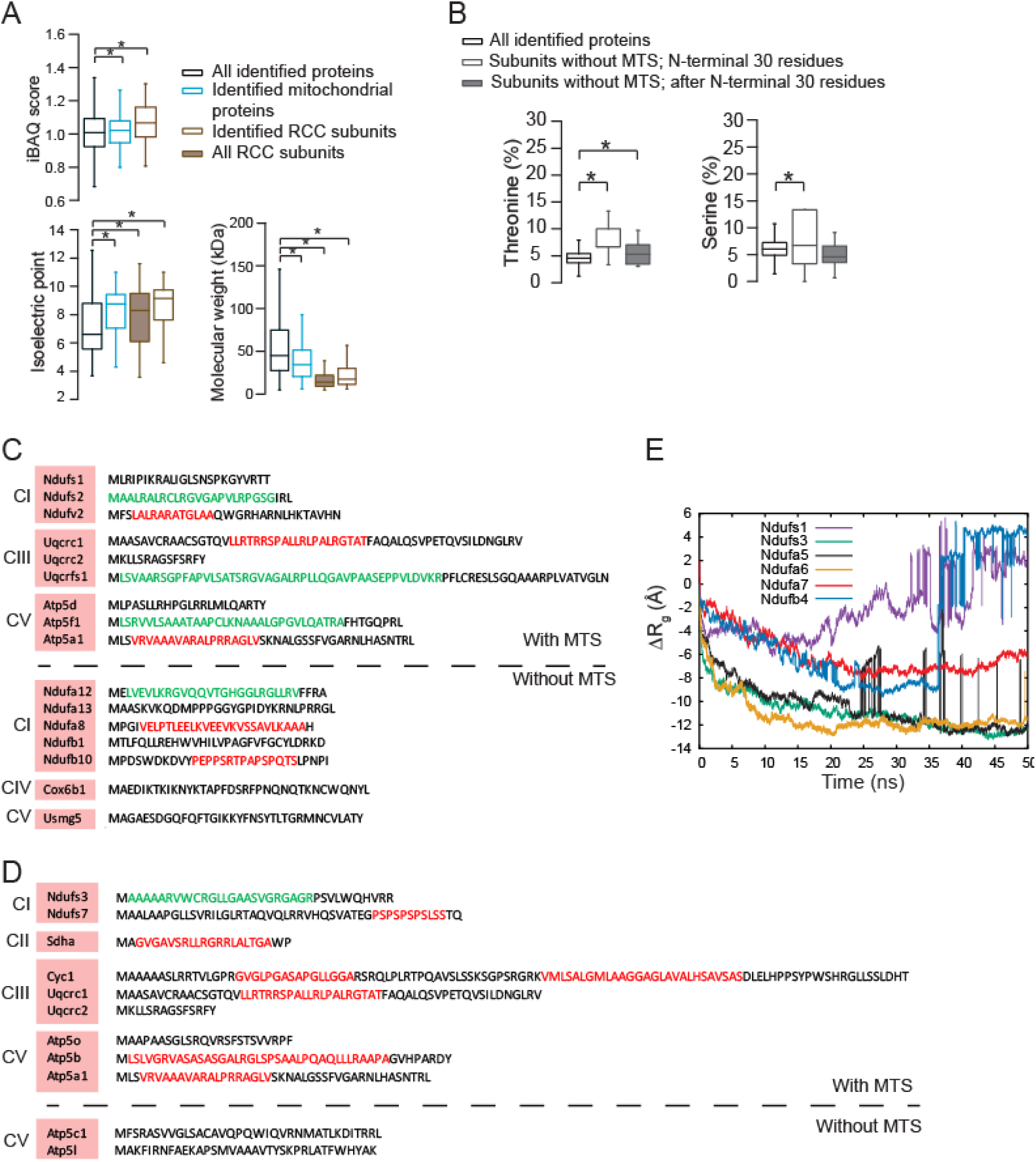
Role of LCR containing N-terminal regions of RCC subunits in aggregation. Related to Figure 4. (A) Box plots showing physicochemical properties of RCC subunits. iBAQ indicates intrinsic abundance of proteins as calculated from mass spectrometry data from the control DMSO treated cells (Note: iBAQ calculation is only possible for identified subunits by mass spectrometry). A separate group encompassing all RCC subunits is also shown for molecular weight and isoelectric point analysis. ^*^ indicates p<0.05 by one-way ANOVA, Bonferroni’s post hoc test. (B) Box plots showing amino acid composition of identified RCC subunits without MTS. N-terminal 30 residues and rest of the protein-sequences are grouped separately. ^*^ indicates p<0.05 by one-way ANOVA, Bonferroni’s post hoc test. (C) and (D) Low complexity regions (LCRs) in the predicted MTS or N-terminal 30 residues of RCC subunits enriched in the total and the insoluble fraction respectively. High confidence LCRs are highlighted in red; low confidence in green. (E) Radius of gyration versus time graph as calculated by Molecular dynamics simulation of N-terminal 30 amino acid residues of RCC subunits.

**Figure S5:**
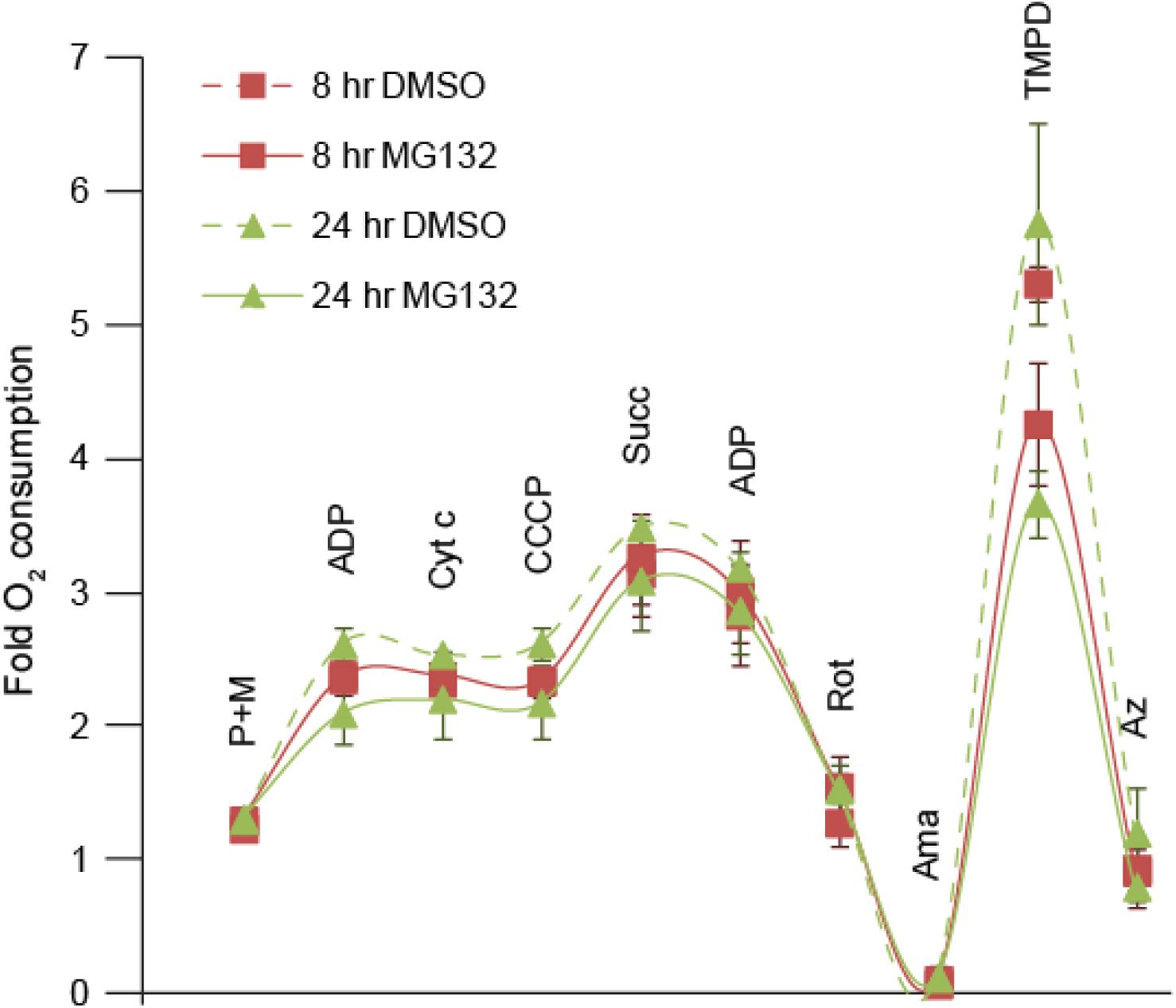
Mitochondrial function in MG132 treated cells - respirometry. Related to Figure 5. Oxygen consumption trace for control (dashed line) and 5 μM MG132 treated (solid line) Neuro2a cells as measured by Oxygraph-2k. Cells were suspended in Mir05 and permeabilized using digitonin followed by addition of respiratory inhibitors and substrates. P+M, pyruvate and malate; ADP, Adenosine diphosphate; Cyt c, Cytochrome c; CCCP, Carbonyl cyanide m-chlorophenyl hydrazone; Succ, Succinate; Rot, Rotenone; Ama, Antimycin A; TMPD, N,N,N’,N’-tetramethyl-p-phenylenediamine; Az, Sodium Azide. SD from 3 independent experiments.

**Figure S6:**
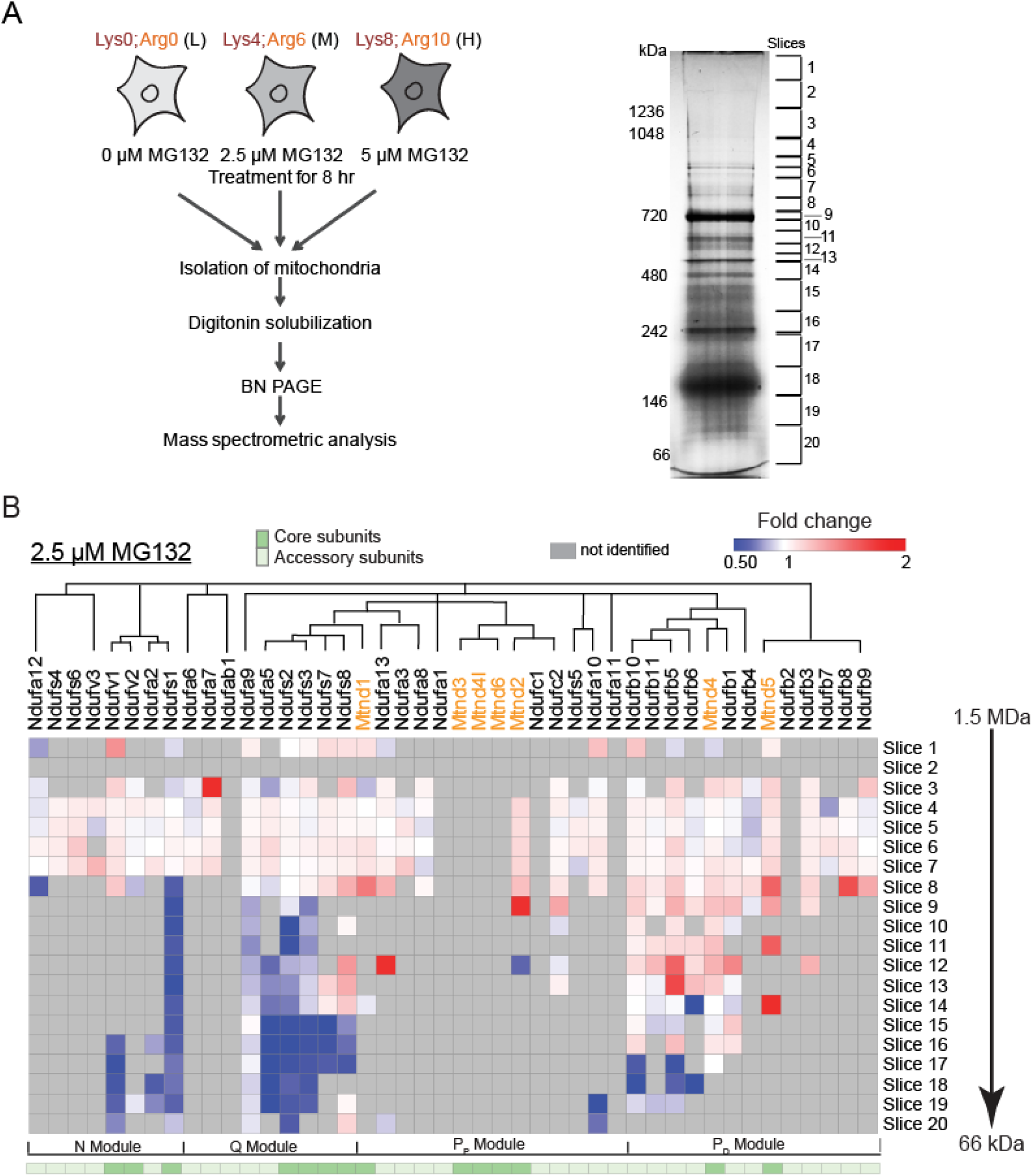
Complexome profiling. Related to Figure 6. (A) Left - schematic describing the SILAC-based quantitative mass spectrometry strategy to profile RCC subunits of MG132 treated Neuro2a cells using BN-PAGE. Right-representative BN-PAGE showing separation of mitochondrial protein complexes. The gel slices are marked and numbered. (B) Alteration of abundance of CI subunits at different slices of BN-PAGE in MG132 treated cells (8 hr) as determined by SILAC-based fold changes over control DMSO treated cells. The subunits are clustered as per CI assembly pathway described by Guerrero-Castillo et al. (Guerrero-Castillo et al., 2017). Mitochondria-encoded subunits are shown in orange.

**Figure S7:**
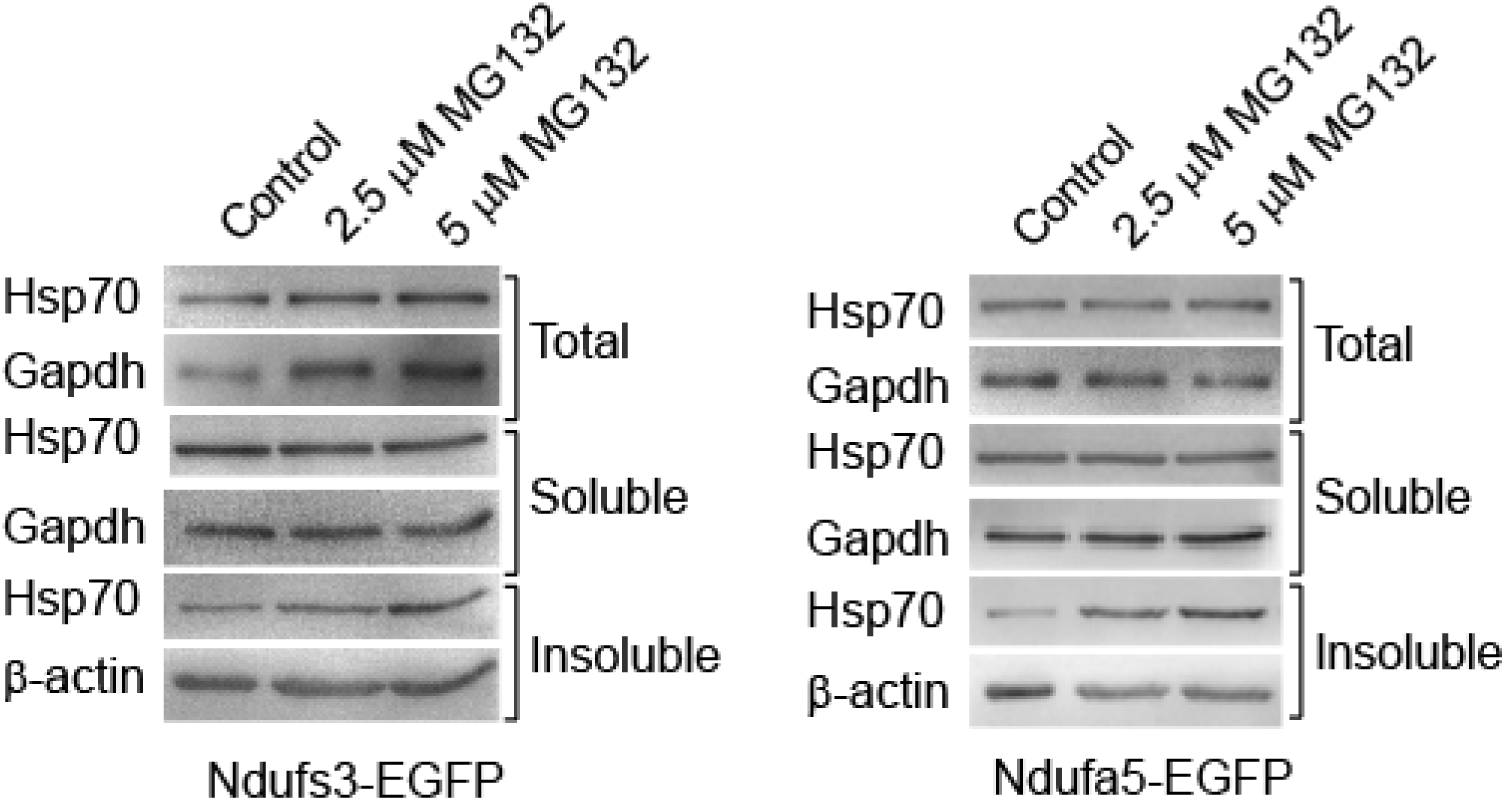
Partitioning of HSP70 to the insoluble fraction of 8 hr MG132 treated Ndufs3-EGFP and Ndufa5-EGFP overexpressing Neuro2a cells. Gapdh served as loading control for total and soluble fraction; β-actin for insoluble fraction.

## Methods

### Constructs

FlucDM-EGFP was PCR-amplified from pCIneo-FlucDM-EGFP (Gupta et al., 2011) and subcloned into pcDNA4/TO using the restriction enzymes KpnI and XbaI (Thermo Scientific). Ndufs3, Ndufa5, Ndufa6 and Ndufb10 were PCR-amplified from Neuro2a cDNA and cloned into pcDNA4/TO EGFP using the restriction enzymes KpnI and XhoI. Snca was PCR-amplified from pRK172/α-synuclein (Ahmad et al., 2006) and cloned into pcDNA4/TO EGFP using KpnI and XhoI. The first 30 residues were removed to prepare the deletion constructs and were tagged before EGFP to prepare NTR constructs. pCI-His-hUbi was from Addgene (Plasmid #31815).

### Cell culture and microscopy

Neuro2a cells were maintained in Dulbecco’s Modified Eagle’s Medium (Gibco) supplemented with 10% fetal bovine serum (FBS) (Gibco) and 90 U/ml penicillin (Sigma) −50 μg/ml streptomycin (Sigma) at 37°C and 5% CO_2_. Transfection of cells was performed with Lipofectamine 3000 reagent (Invitrogen) according to the manufacturer’s protocol. Mitotracker Red CMXRos (Invitrogen) staining was done at a final concentration of 0.5 mM in culture media by incubating cells for 30 min under normal growth conditions. Slides were prepared by counterstaining with DAPI (Sigma) and observed under Zeiss Axioimager Z.1 Microscope (Figure 1D) or Leica TCS sp8 (Figure 4B and 5B).

For SILAC based mass spectrometry - cells were grown in SILAC DMEM (Thermo Scientific) supplemented with 10% dialyzed FBS (Gibco), 90 U/ml penicillin (Sigma) - 50 μg/ml streptomycin (Sigma) and either Light [L-Lysine 2HCl / L-Arginine HCl (Lys0/Arg0)] or Medium [L-Lysine 2HCl (4,4,5,5-D4) / L-Arginine HCl (^13^C_6_) (Lys4/Arg6)] or Heavy [L-Lysine 2HCl (^13^C_6_, ^15^N_2_) / L-Arginine HCl (^13^C_6_, ^15^N_4_) (Lys8/Arg10)] isotopes of lysine and arginine (Thermo Scientific).

### Cell viability and proteasome activity assay

Cell viability was determined using MTT assay (Sigma). Briefly, 10000 cells were plated in 96-well culture plate. After treatment, MTT solution (0.5 mg/ml in growth medium) was added followed by incubation for further 3 hr under normal growth conditions. Medium was removed and formazan crystals were dissolved in DMSO. Absorbance was measured at 570 nm using PerkinElmer EnSpire Multimode plate reader.

Proteasome activity assay was performed using Proteasome-Glo Chymotrypsin-Like Cell-Based Assay kit (Promega) following the protocol described by the manufacturer. Briefly, 10000 cells were plated in 96-well culture plate. After 8 hr of MG132 treatment, pre-mixed assay buffer containing substrate and luciferin detection reagent was added in equal volume to the sample and incubated for 10 min at RT. The supernatant was transferred to optiplate-96 (white; Perkin Elmer) and luminescence recorded using PerkinElmer EnSpire Multimode plate reader.

Cell death was measured using Pierce LDH Cytotoxicity Assay Kit (Thermo Scientific) as per manufacturer’s protocol. Briefly, 10000 cells were plated in 96-well culture format. After 8 hr of MG132 treatment, equal volume of culture media and reaction mix was taken in a fresh plate and incubated for 30 min in dark at RT. Absorbance was recorded at 490 nm in PerkinElmer EnSpire Multimode plate reader.

### Luciferase assay

Activity of FlucDM was estimated using Pierce Firefly Luciferase Glow Assay Kit (Thermo Scientific). Assay was performed as per kit protocol. Neuro2a cells were transfected with pcDNA4/TO FlucDM-EGFP using lipofectamine 3000. After 24 hr, transfected cells were plated in 96-well culture plate and upon attachment were treated with 2.5 μM and 5 μM MG132 for 8 hr and DMSO as solvent control. The cells were lysed in 100 μl of 1X Lysis buffer provided in the kit and 20 μl of this lysate was transferred into optiplate-96 (white; Perkin Elmer) with 50 μl of working solution containing Luciferin. Luminescence was recorded in PerkinElmer EnSpire Multimode plate reader.

### Western blotting

Cell pellet was lysed in NP-40 lysis buffer (50 mM Tris-HCl [pH 7.8], 150 mM NaCl, 1% NP-40, 0.25% Sodium deoxycholate, 1 mM EDTA, protease inhibitor cocktail (Roche)) at 4°C for 45 min with intermittent vortexing. Lysed cells were centrifuged at 12000g for 15 min at 4°C and the supernatant was collected as the soluble fraction. The remaining pellet was washed twice with 1X PBS and boiled in 4X SDS loading buffer (0.2 M Tris-HCl [pH 6.8], 8% SDS, 0.05 M EDTA, 4% 2-mercaptoethanol, 40% glycerol, 0.8% Bromophenol blue) for 15 min to get insoluble fraction. Total fraction was prepared by directly dissolving the cell pellet in 4X SDS loading buffer for 15 min. Protein fractions were separated by SDS-PAGE and transferred onto 0.2 μm PVDF membrane (Bio-rad) for 90 min (for histones 60 min) at 300 mA using the Mini-Trans Blot cell system (Bio-rad). Membranes were probed by appropriate primary and secondary antibodies (**Supplementary Methods Table 1**) and imaged using documentation system (Vilber Lourmat).

### Reverse Transcriptase PCR (RT-PCR)

Total RNA was prepared using Trizol method (Ambion) according to the manufacturer’s protocol and treated with DNase I (Ambion). RNA concentration was measured using Nanodrop2000 spectrophotometer (Thermo Scientific). For cDNA synthesis, 2 μg of total RNA was used along with SuperScript III Reverse Transcriptase (Invitrogen) in a final volume of 20 μl according to the kit protocol. Quantitative PCR was carried out in Applied Biosystems 7900HT Fast Real-Time PCR System using Power SYBR green Master-mix (Applied Biosystems). (**Supplementary Methods Table 3**)

### Sample preparation for mass spectrometry

1.5 million cells were plated in 100 mm culture dishes. The light-labeled cells (L) served as solvent control whereas medium-labeled (M) and heavy-labeled (H) cells were treated with 2.5 μM and 5 μM of MG132 respectively. After 8 hr of incubation, equal number of L-, M- and H-labeled cells were pooled together and total, soluble and insoluble fractions were prepared as described earlier. The fractions were separated on NuPAGE 4-12% Bis-Tris Protein Gels (Invitrogen). The gel was run in MES buffer (100 mM MES, 100 mM Tris-HCl, 2 mM EDTA, 7 mM SDS) at 200V for 40 min, fixed and stained with coomassie brilliant blue. Preparation of gel slices, reduction, alkylation, and in-gel protein digestion was carried out as described by (Shevchenko et al., 1996). Finally, peptides were desalted and enriched according to (Rappsilber et al., 2003).

### LC-MS/MS

Peptides eluted from desalting tips were dissolved in 2% formic acid and sonicated for 5 min. Soluble fraction was analyzed on Linear Trap Quadrupole (LTQ)-OrbitrapVelos interfaced with nanoflow LC system (Easy nLC II, Thermo Scientific). Peptide fractions were separated on a Bio Basic C18 pico-Frit nanocapillary column (75 μm × 10 cm; 3 μm) using a 120 min linear gradient of the mobile phase [5% ACN containing 0.2% formic acid (buffer-A) and 95% ACN containing 0.2% formic acid (buffer-B)] at a flow rate of 300 nL/min. Full scan MS spectra (from *m*/*z* 400–2000) were acquired followed by MS/MS scans of Top 20 peptides with charge states 2 or higher.

Total and insoluble fractions were analyzed on Q Exactive (Thermo Scientific) interfaced with nanoflow LC system (Easy nLC II, Thermo Scientific). Peptide fractions were separated on a Bio Basic C18 pico-Frit nanocapillary column (75 μm × 10 cm; 3 μm) using a 60 min linear gradient of the mobile phase [5% ACN containing 0.2% formic acid (buffer-A) and 95% ACN containing 0.2% formic acid (buffer-B)] at a flow rate of 400 nL/min. Full scan MS spectra (from *m*/*z* 400–2000) were acquired followed by MS/MS scans of Top 10 peptides with charge states 2 or higher.

### Peptide identification and statistical analysis

For peptide identification, raw MS data files were loaded onto MaxQuant proteomics computational platform (Ver. 1.3.0.5) (Cox and Mann, 2008) and searched against Swissprot database of *Mus musculus* (release 2016.03 with 16790 entries) and a database of known contaminants. MaxQuant used a decoy version of the specified database to adjust the false discovery rates for proteins and peptides below 1%. The search parameters included constant modification of cysteine by carbamidomethylation, enzyme specificity trypsin, multiplicity set to 3 with Lys4 and Arg6 as medium label and Lys8 and Arg10 as heavy label. Other parameters included minimum peptide for identification 2, minimum ratio count 2, re-quantify option selected and match between runs with 2 min time window. iBAQ (Schwanhausser et al., 2011) option was selected to compute abundance of the proteins. Bioinformatics and statistical analysis was performed in Perseus environment (Ver. 1.5.2.4) (Tyanova et al., 2016).

For total and soluble fractions, M/L and H/L ratios were converted into log2 space, and average ratios and SD (standard deviations) were calculated for each data set. The log2 M/L and H/L ratio of each protein were converted into a z-score, using the following formula:

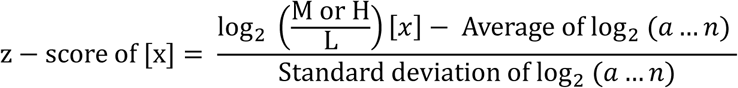

where x is a single protein in the data set population (a….n). The z-score was a measure of how many Standard deviation (σ) units, the log2 M/L or H/L ratio of the protein was away from the population mean. A z-score ≥ 1.96σ represented that differential expression of the protein lied outside the 95% confidence interval and were considered to be significant. We divided the protein population into 5 divisions according to their z-score distribution. Proteins having (−1.96≥z ≥1.96) were termed as Highly depleted and Highly enriched, respectively. The proteins with (1.96>z>1) or (−1.96<z<-1) were classified as Enriched and Depleted respectively. The proteins with (1≥z≥-1) scores were grouped as Unchanged. As insoluble fraction showed skewed distribution, mean and standard deviation of total fraction was used to normalize and calculate z-score.

### Gene ontology and bioinformatics analysis

Gene ontology (GO) analysis was performed using GOrilla - Gene Ontology enRIchment anaLysis and visuaLizAtion tool (Eden et al., 2009). All proteins identified in the mass spectrometry experiments across the fractions were used to prepare the background set with 10^-4^ as p-value threshold. Further, Revigo was used for more stringent representation of GO analysis data (small-0.5) (Supek et al., 2011). Subcellular localization of the proteins was acquired from UniProt (Apweiler et al., 2004). Protein abundance was calculated by selecting iBAQ option in MaxQuant tool. iBAQ values for the DMSO control (not MG132 treated) were used to calculate the relative abundances under normal conditions for the proteins identified and converted into log space. Camsol was used to predict instrinsic solubility based on the protein sequence (Sormanni et al., 2015). The molecular weight and other properties of the identified proteins (isoelectric point (pI), amino acid composition, hydropathicity etc.) were taken from ProtParam (ExPASy web server) (Gasteiger et al., 2005). TMHMM Server v. 2.0 was used to predict transmembrane helices (Krogh et al., 2001). Low complex regions in proteins were predicted by SEG-prediction of low complex region (Wootton, 1994). For prediction of mitochondrial target sequence, TPpred 2.0 online tool was used (Savojardo et al., 2014).

### Secreted protein analysis

Cells were transfected with EGFP tagged synuclein for 24 hr. Culture media was replaced with fresh DMEM before 2.5 μM and 5 μM MG132 treatment to the cells. Media was collected post 8 hr and 24 hr from treated cells and subjected to centrifugation at 10000g for 30 min at 4°C. For immunoblotting, equal volume of each sample was loaded after boiling with SDS loading buffer for 10min and blotted using anti-EGFP.

### His-ubiquitin pull down assay

Cells transfected with EGFP tagged subunits and pCI-His-hUbi were treated with 5 μM MG132 for 8 hr. Cells were washed with ice-cold PBS and lysed in lysis buffer (6 M Guanidine-HCl, 0.1 M HEPES [pH 7.4], 5 mM Imidazole). Ni-NTA agarose beads (Invitrogen) were equilibrated with lysis buffer and added to each cell lysate and incubated at room temperature for 2 hr in a rotating wheel. Beads were washed twice with lysis buffer and four times with wash buffer (300 mM NaCl, 50 mM Tris-HCl [pH 7.4], 20 mM imidazole, 1% NP-40). Elution was performed by boiling the beads in 2x SDS PAGE loading buffer supplemented with 300 mM imidazole for 15 min. Samples were separated on SDS-PAGE and immunoblotting was done using anti-EGFP.

### Molecular Dynamics Simulation

All-atom, explicit solvent molecular dynamics simulation was used to study the aggregation propensity of the first 30 amino acids of Ndufs3, Ndufa5, Ndufa6, Ndufs1, Ndufa7 and Ndufb4. Molecular dynamics simulation software CHARMM (Chemistry at HARvard Macromolecular Mechanics) (Brooks et al., 2009) and NAMD (NAno scale Molecular Dynamics) (Phillips et al., 2005) with CHARMM36 force field (Best et al., 2012) were used to model and simulate the peptides. Tesla K40 GPU card was used for faster simulation process. Extended form of each peptide was used as the starting conformation. Each peptide-segment was then copied into nine other similar segments using CHARMM and translated in space randomly, resulting in 10 copies of the peptides. Each system, containing 10 peptide-segments, was energy minimized and solvated in an orthogonal water box having dimension x=162, y=83, z=75 Å using CHARMM_GUI webserver (Jo et al., 2008). The system was then heated to 310K and simulated for 50 ns using NAMD. SHAKE algorithm has been used to fix the hydrogen bond distances during the simulation (Ryckaert, 1977) Integration time step was taken to be 1 fs and cutoff was set at 12.0 Å to adjust both electrostatic and van der Waals interactions.

### High-resolution respirometry

For individual respiratory complex activities, oxygen consumption was evaluated using the substrate-uncoupler-inhibitor titration reference protocol (Pesta and Gnaiger, 2012) with Oxygraph-2k (Oroboros Instruments, Austria). Neuro2a cells (1X10^6^ cells) were suspended in mitochondrial respiration medium Mir05 (110 mM Sucrose, 0.5 mM EGTA, 3.0 mM MgCl_2_, 80 mM KCl, 60 mM K-lactobionate, 10 mM KH_2_PO_4_, 20 mM Taurine, 20 mM Hepes, 1.0 g/l BSA, pH 7.1). Cells were permeabilized using digitonin (5 μg/ml) followed by respiratory complex inhibitors and substrates. Cytochrome c test was done to check the intactness of outer mitochondrial membrane. Oxidative phosphorylation capacity by Complex I was determined by the addition of 5 mM Pyruvate and 5 mM Malate followed by the addition of 2.5 mM ADP. Complex II oxidative phosphorylation was measured after addition of 0.5 μM Rotenone and 10 mM Succinate. Non-mitochondrial respiration was observed by addition of 2.5 μM Antimycin A. For complex IV assay, 2 mM Ascorbate and 0.5 mM N,N,N’,N’-tetramethyl-p-phenylenediamine (TMPD) were added followed by 200 mM Sodium azide. All reagents were procured from Sigma.

ROS levels were measured with Cellular ROS/Superoxide Detection Assay Kit (Abcam) as per manufacturer’s protocol. Briefly, treated cells were detached and incubated with ROS Detection solution for 30 min at 37°C in dark. Samples were analysed using Gallios Flow Cytometer (Beckman Coulter).

### Mitochondria isolation

Mitochondria isolation from cultured cell lines was performed according to Schägger (1995) and Rebeca Acín-Pérez (2008), with some modifications (Pfeiffer et al., 2003; Schagger, 1995). Briefly, MG132 treated L-, M- and H-labeled cells as mentioned earlier were pooled together (10 million) and pellet was washed twice with PBS. Cell pellet was frozen at −80°C to increase cell breakage and homogenized in a dounce homogenizer with about 10 cell pellet volumes of homogenizing buffer, buffer A (83 mM Sucrose, 10 mM MOPS [pH 7.2]). After adding an equal volume of buffer B (250 mM Sucrose, 30 mM MOPS [pH 7.2]), nuclei and unbroken cells were removed by centrifugation at 1000g during 5 min at 4°C. Mitochondria were collected from the supernatant by centrifuging at 12000g during 5 min and washed twice under the same conditions with buffer C (320 mM Sucrose, 1 mM EDTA, 10 mM Tris-HCl [pH 7.4]). Mitochondrial pellets were resuspended in buffer D (1 M 6-aminohexanoic acid, 50 mM Bis-Tris HCl [pH 7.0]), and estimated by Pierce Coomassie Plus (Bradford) Assay Reagent (Thermo Scientific).

### BN-PAGE and Complexome profiling

For complexome profiling 100-120 μg mitochondria were solubilized by adding 8 g digitonin (Sigma) per g of mitochondria, and incubating 5 min in ice. After a 30 min centrifugation at 13000g, the supernatant was collected, and one-third of the final volume of the sample, 5% coomassie brilliant blue G-250 dye in 1 M 6-aminohexanoic acid was added. Glycerol was added at final concentration of 10% and the sample was loaded onto NativePAGE 3-12% Bis-Tris Protein Gels (Invitrogen). Native PAGE was run using anode buffer ( 50 mM Bis-Tris HCl [pH 7.0]) and blue cathode buffer (50 mM Tricine, 15 mM Bis-Tris HCl [7.0], 0.02% coomassie brilliant blue) at 100 V until samples entered separating gel, then run at 150 V until dye-front reached 1/3 the length of the entire gel. At this point blue cathode buffer was replaced with dye-free cathode buffer for remainder of run at 200 V until dye-front reached end of gel. The gel was fixed and stained with coomassie brilliant blue and cut into 20 slices for mass spectrometry. Preparation of gel slices, reduction, alkylation, and in-gel protein digestion was carried out as described by (Shevchenko et al., 1996). Finally, peptides were desalted and enriched according to (Rappsilber et al., 2003). Peptides eluted from desalting tips were dissolved in 2% formic acid and sonicated for 5 min. Samples were analysed on Q-Exactive HF (Thermo Scientific) and were separated on a EASY-Spray PepMap RSLC C18 Column (75 μm × 15 cm; 3 μm) using a 60 min linear gradient of the mobile phase [5% ACN containing 0.2% formic acid (buffer-A) and 95% ACN containing 0.2% formic acid (buffer-B)] at a flow rate of 300 nL/min. Full scan MS spectra (from *m*/*z* 400–1650) were acquired followed by MS/MS scans of Top 10 peptides with charge states 2 or higher. Raw MS data files of individual slices were loaded and analysed separately using MaxQuant (Ver. 1.3.0.5) (Cox and Mann, 2008) and searched against Swissprot database of *Mus musculus* (release 2016.03 with 16790 entries) and a database of known contaminants.

---

**Supplementary Methods Table 1.** Primary Antibody list:

Primary Antibody list:
| Antibody Name | Company | Catalog number | Dilution |
| --- | --- | --- | --- |
| Mouse monoclonal Ubiquitin (P4D1) | Santa cruz biotechnology | Cat#sc-8017 | 1:1000 |
| Mouse monoclonal Anti-GAPDH [6C5] antibody | abcam | Cat#ab8245 | 1:10000 |
| Rat monoclonal Anti-HSF1 antibody [10H8] | abcam | Cat#ab61382 | 1:1000 |
| Mouse monoclonal Anti-HSP70/HSC70 (N27F3-4) | Enzo life sciences | Cat#ADI-SPA-820-F | 1:1000 |
| Rabbit polyclonal Anti-LC3B antibody | abcam | Cat#ab51520 | 1:3000 |
| Rabbit polyclonal Anti-GRP78 BiP antibody | abcam | Cat#ab21685 | 1:1000 |
| Mouse monoclonal Anti-Hsp60 antibody [LK-1] | abcam | Cat#ab59457 | 1:1000 |
| Rabbit monoclonal Anti-VDAC1/Porin antibody [EPR10852(B)] - Mitochondrial Loading Control | abcam | Cat#ab154856 | 1:10000 |
| Rabbit polyclonal Anti- | abcam | Cat#ab1791 | 1:2000 |
| Histone H3 - Nuclear<br>Loading Control and ChIP<br>Grade |  |  |  |
| Chicken polyclonal Anti-GFP antibody | abcam | Cat#ab13970 | 1:5000 |
| Mouse monoclonal Anti-c-Myc antibody [9E10] - ChIP<br>Grade (ab32) | abcam | Cat#ab32 | 1:1000 |
| Rabbit polyclonal Anti-beta<br>Tubulin - Loading control | abcam | Cat#ab6046 | 1:1000 |
| Mouse monoclonal Anti-beta<br>Actin antibody - Loading<br>control | abcam | Cat#ab8224 | 1:10000 |
| Mouse monoclonal Anti-Histone H3 (tri methyl K4)<br>antibody - ChIP Grade | abcam | Cat#ab1012 | 1:1000 |
| Goat polyclonal Anti-Histone<br>H3 (di methyl K4) antibody -<br>ChIP Grade | abcam | Cat#ab11946 | 1:1000 |
| Mouse monoclonal Anti-Histone H3 (tri methyl K27)<br>antibody - ChIP Grade | abcam | Cat#ab6002 | 1:1000 |
| Anti-Histone H3 (tri methyl<br>K36) antibody - ChIP Grade | abcam | Cat#ab9050 | 1:1000 |
| Anti-Histone H3 (acetyl | abcam | Cat#ab4729 | 1:1000 |
| K27) antibody - ChIP Grade |  |  |  |
| Rabbit polyclonal Anti-Histone H4 (tri methyl K20) antibody - ChIP Grade | abcam | Cat#ab9053 | 1:1000 |
| Rabbit monoclonal Anti-Histone H4 (acetyl K5+K8+K12+K16) antibody [EPR16606] | abcam | Cat#ab177790 | 1:1000 |
| Rabbit Anti-Rat IgG H&L (HRP) | abcam | Cat#ab6734 | 1:5000 |
| Anti-Rabbit IgG (Whole molecule) (peroxidase conjugate) | sigma | Cat#A6154 | 1:20000 |
| Anti-mouse IgG (whole molecule) (peroxidase conjugate) | sigma | Cat#A4416 | 1:20000 |
| Goat Anti-Chicken IgY H&L (HRP) | abcam | Cat#ab205721 | 1:10000 |
| Rabbit Anti-Goat IgG H&L (HRP) | abcam | Cat#ab6741 | 1:20000 |

**Supplementary Methods Table 2.**
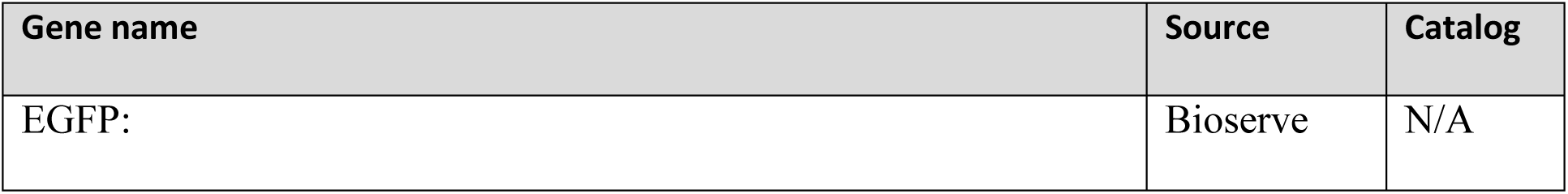
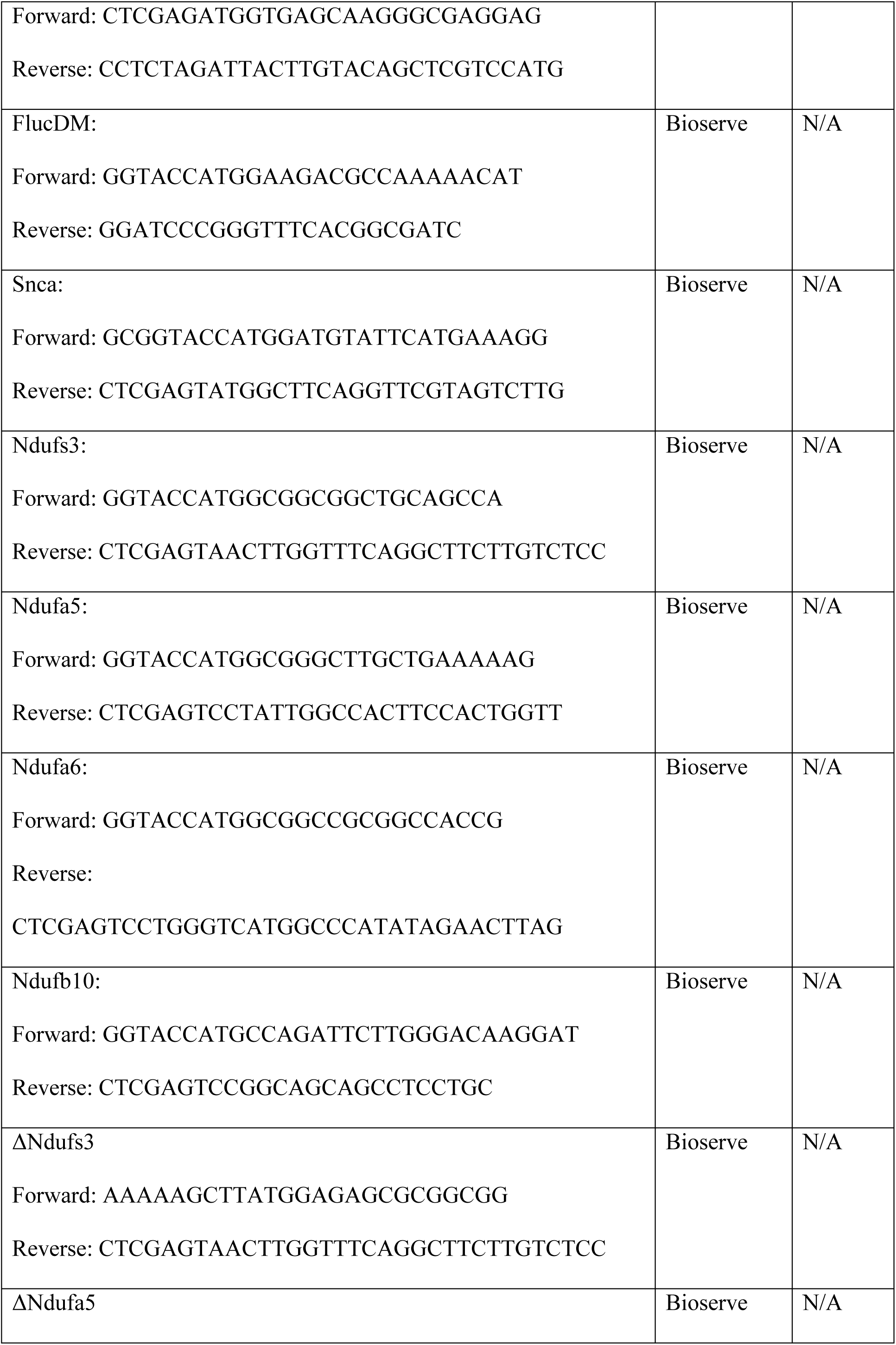
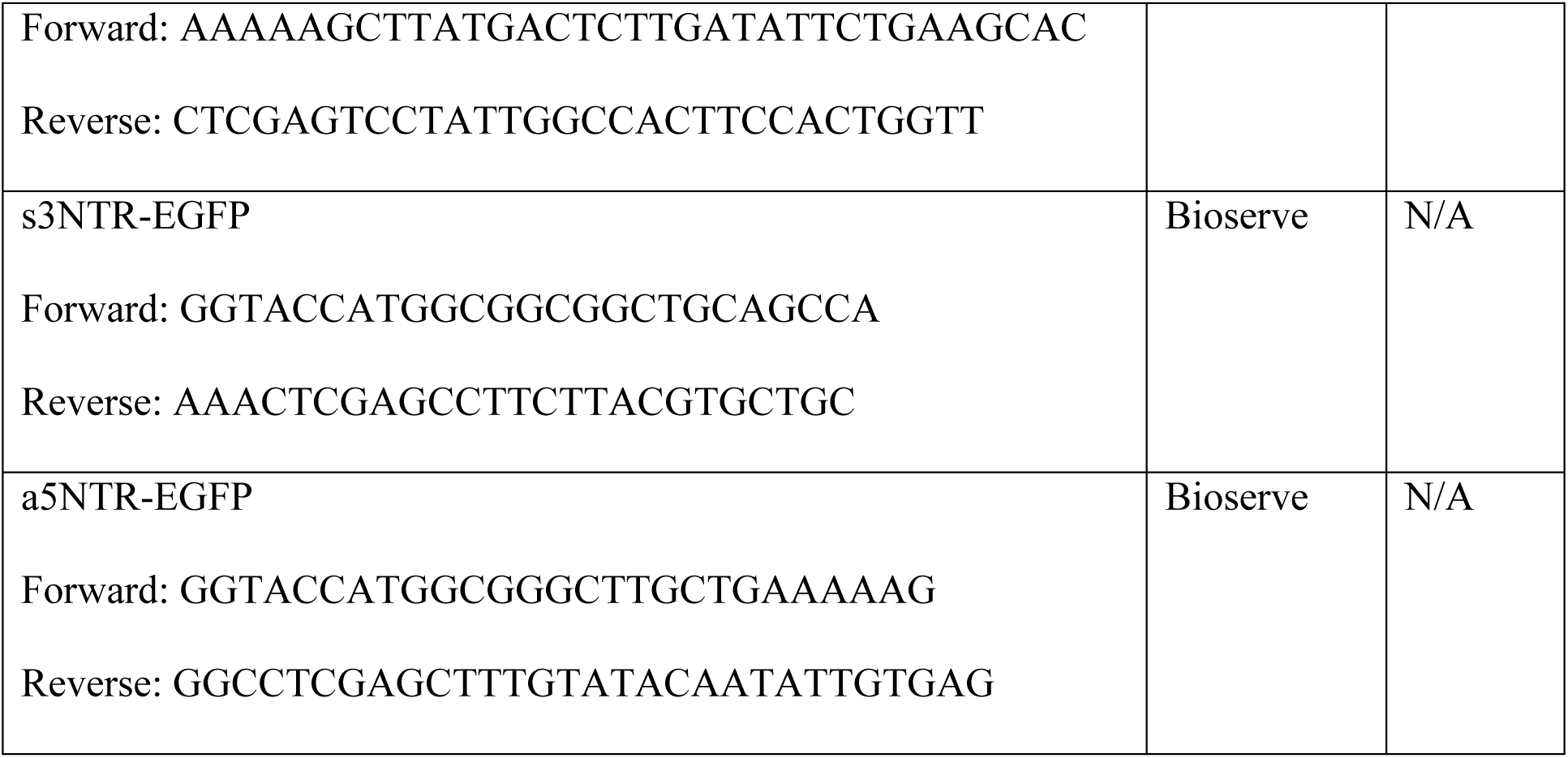
Primers for Cloning:

**Supplementary Methods Table 3.**
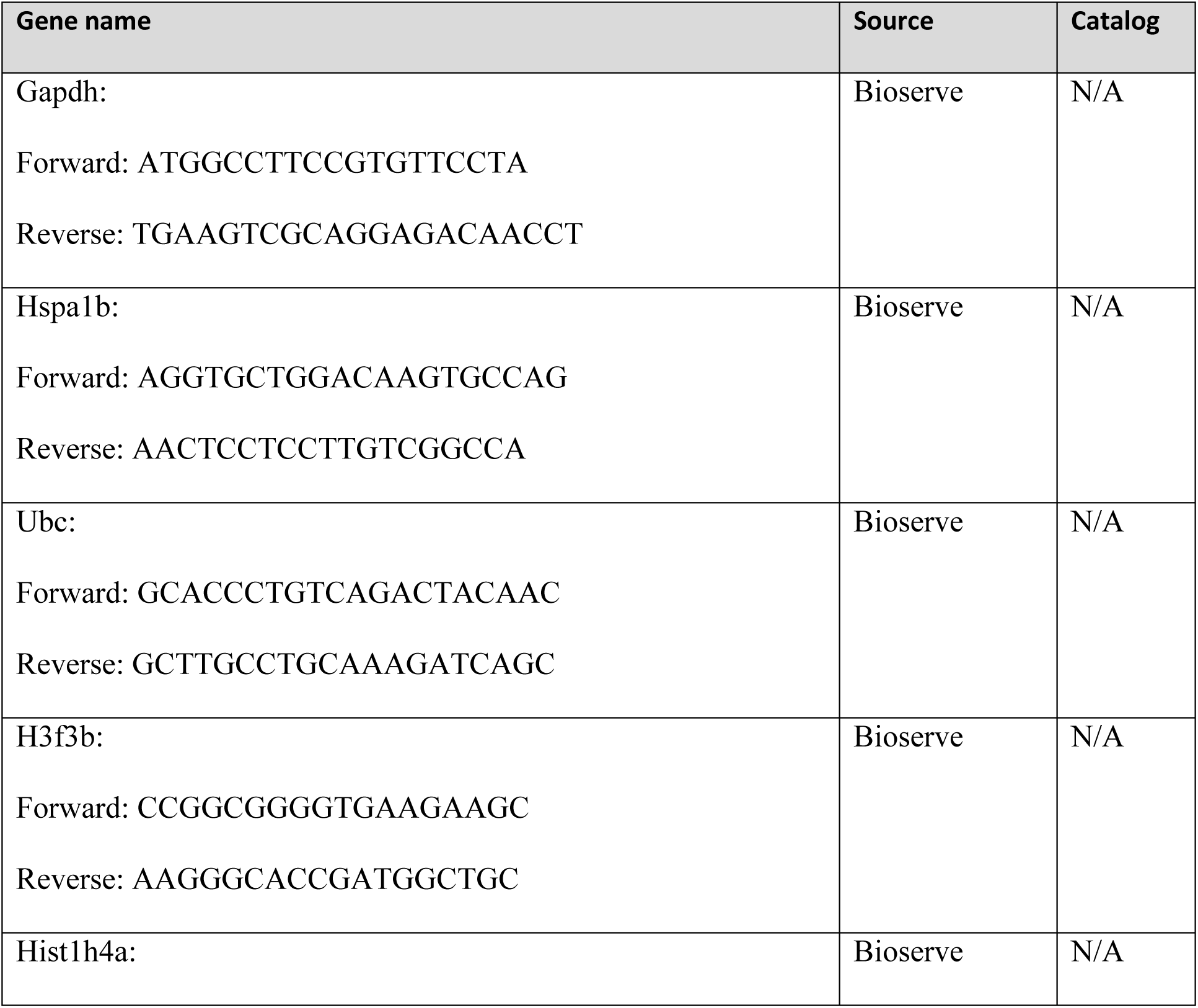
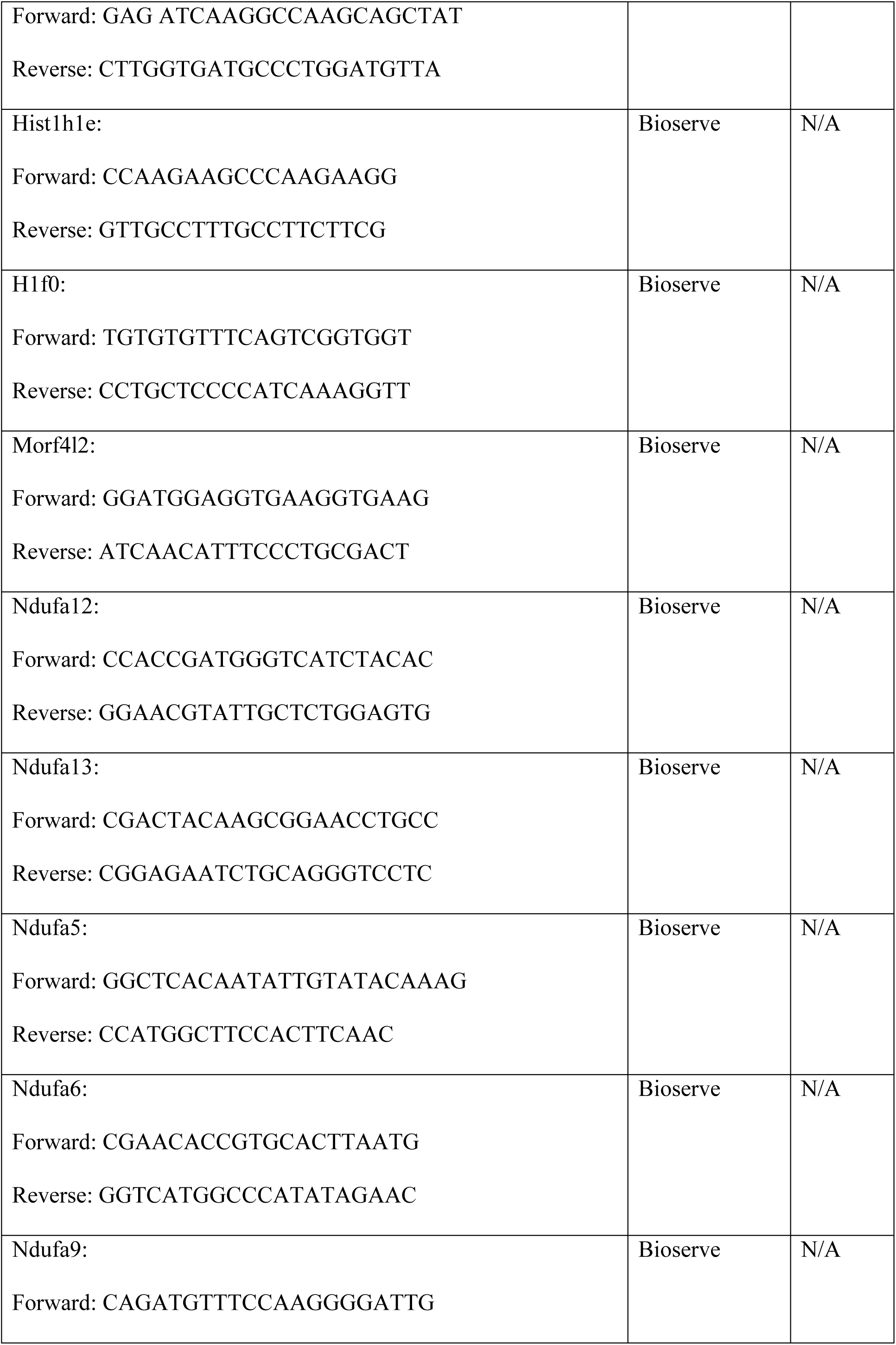
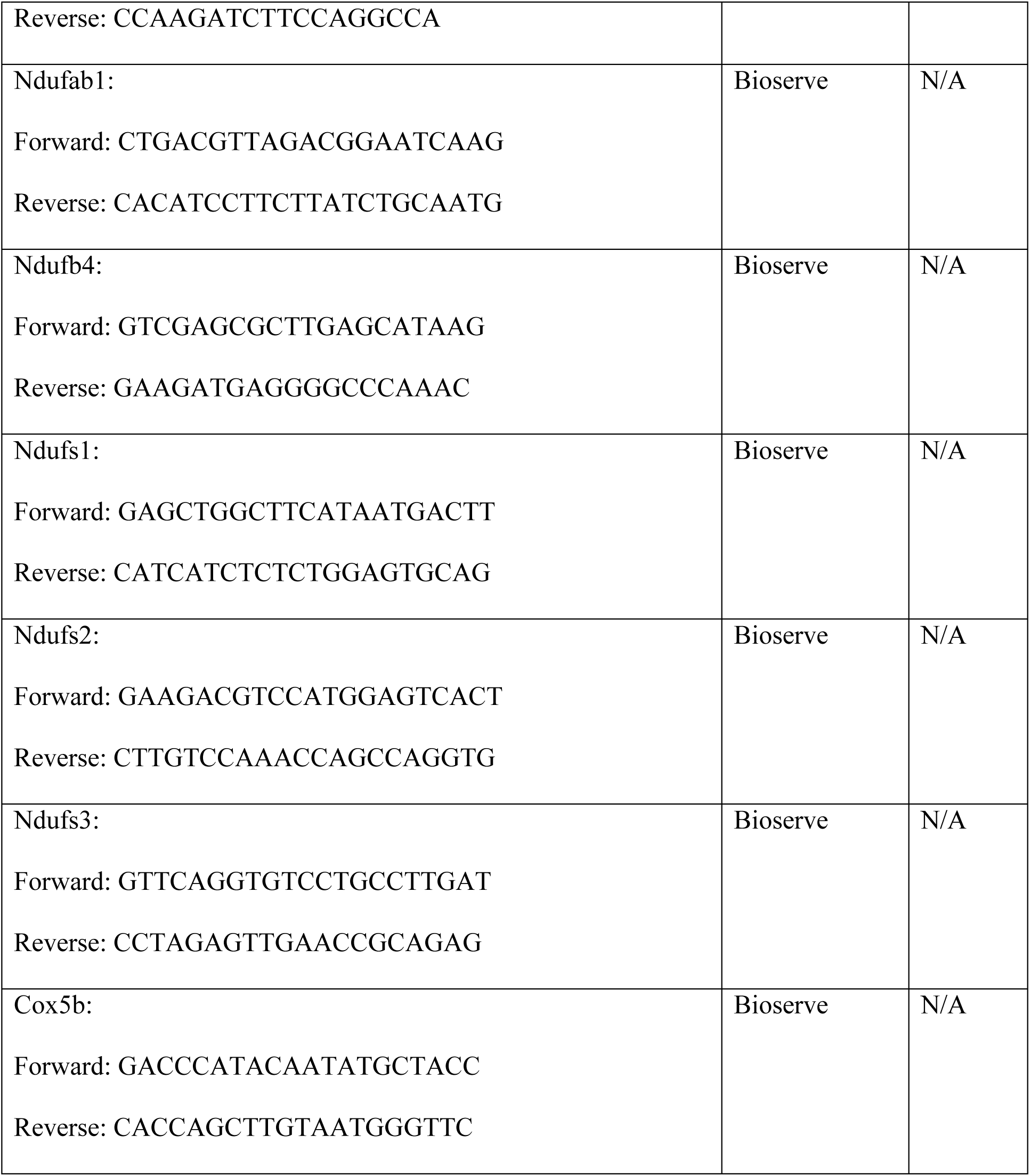
Primers for RT-PCR:

